# Distinct developmental trajectories of externally- and internally-generated hippocampal sequences and assemblies

**DOI:** 10.1101/2025.11.27.690943

**Authors:** Erwan Leprince, Caroline Filippi, Manon Mantez, Robin F. Dard, Vito Dichio, Jure Majnik, Marina Cretella, Marco Bocchio, Michel A. Picardo, Rémi Monasson, Jean-Claude Platel, Rosa Cossart

**Affiliations:** INMED, INSERM, Aix-Marseille University, Turing Centre for Living systems, Marseille, France; Department of Physiology, Development and Neuroscience, University of Cambridge, Cambridge, United Kingdom; Laboratory of Sensory Processing, Brain Mind Institute, School of Life Sciences, Ecole Polytechnique Fédérale de Lausanne (EPFL), Lausanne, Switzerland; Laboratoire de Physique de l’ENS, PSL & CNRS UMR8023, Sorbonne Université, Paris, France; Department of Psychology, Durham University, Durham, United Kingdom; Learning and Memory Processes Centre, Durham University, Durham, United Kingdom

## Abstract

The hippocampus generates cognitive maps through two distinct mechanisms: external representations anchored to environmental landmarks and internal sequences integrating self-referenced information. In adult CA1, these functions segregate radially, with deep cells processing external cues and superficial cells handling internal representations. This anatomical organization suggests that developmental processes constrain the formation of these functionally distinct circuits. However, the developmental timeline for the emergence of these two types of hippocampal representations remains unknown. Using longitudinal two-photon calcium imaging in head-fixed mice during the third postnatal week, we investigated when internal versus external hippocampal representations emerge. We recorded from deep and superficial CA1 layers while mice ran on treadmills that are bare or enriched with tactile cues, addressing the challenge of tracking cells across sessions. Internal sequences emerge during the mid-fourth postnatal week, following the development of cue-based representations. These sequences initially encode elapsed time before evolving to integrate run distance, arising after rest cell assemblies as spatial coding and functional connectivity networks stabilize. These findings identify the mid fourth postnatal week as a critical developmental milestone marking the establishment of a balanced integration between external landmark-based and internal self-referenced spatial representations. This transition period may constitute a vulnerable developmental window during which disruptions could predispose to disorders characterised by altered cognitive maps.

**Highlights:**

- P24 is critical for CA1 development
- Cue-based representations stabilize by P24
- Internal sequences emerge at P24, encoding run duration before distance
- Cell assemblies form before internal sequences

## Introduction

The hippocampus supports multiple cognitive functions, including spatial navigation and memory formation ^1–3^. Converging evidence indicates that the generation of sequential patterns of cell assembly activation is critical to its role in cognition. These hippocampal sequences organise experience into structured representations—or *cognitive maps*—that encode not only spatial relationships between locations but also temporal, social, and abstract dimensions of experience, such as elapsed time or sensory variables ^3–5^. Hippocampal sequences depend on two distinct mechanisms: one that produces representations anchored to external environmental landmarks, and another that structures neuronal activity into internal sequences integrating self-referenced information independently of physical location, such as run distance or elapsed time ^1,6–12^. These sequences can unfold across different timescales, from subsecond intervals to behavioral timescales spanning several seconds. Behavioral-timescale sequences bind together cell assemblies activated during synchronous network events occurring during rest^13^.

In the adult CA1 region, these two forms of representation segregate into distinct subcircuits along the radial axis. Deep CA1 principal cells are predominantly anchored to external sensory cues (external representation), while superficial cells convey self-referenced and predictive information (internal representation)^6,7,14,15^. Since this radial axis reflects a developmental gradient of neurogenesis, this functional segregation suggests that the organization of hippocampal CA1 circuits could be established by early developmental events ^16^.

Early postnatal hippocampal activity is initially anchored to extrinsic sensorimotor inputs, such as twitches and spontaneous movements, which drive correlated neuronal activity during the first and second postnatal weeks ^17–20^. These early activity patterns, supported by the emergence of local recurrent connectivity ^17,19^, may provide a substrate for the formation of structured neuronal sequences. By the third postnatal week, sequences of place cells can already be detected in cue-rich environments ^21–24^, reflecting externally driven spatial representations. Around this same period, hippocampal circuits also begin exhibiting compressed theta sequences, which combine both external and self-referenced information ^23,25^.

Despite detailed characterisation of early place cell and theta sequence emergence, the relative timing between these internal and external representations during postnatal development remain unresolved. Thus far, evidence appears mixed. In rodents, the head-direction system—a self-referenced framework for spatial orientation—emerges before the formation of stable place cells, which are anchored to environmental landmarks ^26^. However, this arguably represents a simpler form of self-referenced coding than the internally generated sequential activity that signals travelled distance or elapsed time ^3,9,11^. In humans, developmental evidence is similarly ambiguous: allocentric navigation strategies generally emerge later than egocentric ones ^27^, suggesting that self-referenced coding may precede landmark-based representations; yet young children can use landmark cues for navigation but fail to integrate them with self-motion information ^28^. These apparently conflicting findings raise a fundamental question of directionality: whether self-referenced representations scaffold the formation of external spatial maps or, conversely, whether external landmarks provide the initial framework upon which internal sequences are constructed.

Our study leverages a unique paradigm capable of isolating self-referenced activity ^11^, providing an opportunity to directly address this fundamental question. Here, we examined the developmental timelines for the emergence of internal and external hippocampal representations in deep and superficial CA1 layers using longitudinal two-photon calcium imaging. Head-fixed mice were free to run or rest on a self-paced treadmill that was either bare or enriched with tactile cues during the fourth postnatal week, allowing us to track neuronal activity across days using a robust cell tracking algorithm ^29^.

Our aims were to determine: (1) when and how the hippocampus begins forming self-referenced neuronal sequences and assemblies; (2) how their developmental trajectories differ from cue-driven representations; and (3) the relative influence of age and experience in this process.

We find that internal sequences emerge later than cue-based spatial representations, marking a developmental transition in CA1 functional organisation during the fourth postnatal week. These results identify a critical period during which the hippocampal network shifts from externally anchored to internally generated dynamics, suggesting early developmental origins for the mechanisms supporting trajectory representation and predictive coding in the mature hippocampus.

## Results

### Cue-based CA1 representations emerge early and stabilize by mid-fourth postnatal week

In order to describe the developmental evolution of CA1 hippocampal dynamics *in vivo*, we expressed the genetically-encoded calcium indicator GCaMP6s in both pyramidal cells and interneurons of the CA1 hippocampus using intraventricular viral infection at birth (Fig. 1A, see Methods). A chronic glass window was implanted at P15 or P16 just above the dorsal hippocampus to image neuronal calcium dynamics. Following 3 to 4 days of recovery and habituation, longitudinal imaging of the calcium fluorescence signal from CA1 pyramidal layer GCaMP6s-expressing neurons was performed every day in mice free to run on a self-paced treadmill. Mice were imaged every day during the 4th postnatal week (P20-27, N = 20 mice; Figure 1A-B and methods). Since distinct spatial/non-spatial representations have been reported along the radial axes of the hippocampus ^8,15,16,30–32^, we have simultaneously imaged two sublayers of the CA1 pyramidal layer separated by 30 μm along the radial axis, a deep one closer to the *stratum oriens* and a superficial one closer to the *stratum radiatum* (Figure 1B). Each field of view (FOV) spanned a 400×400 µm² region and enabled the simultaneous imaging of an average of 506 neurons (ranges: 81-1126, from 20 mice).

**Figure 1.**
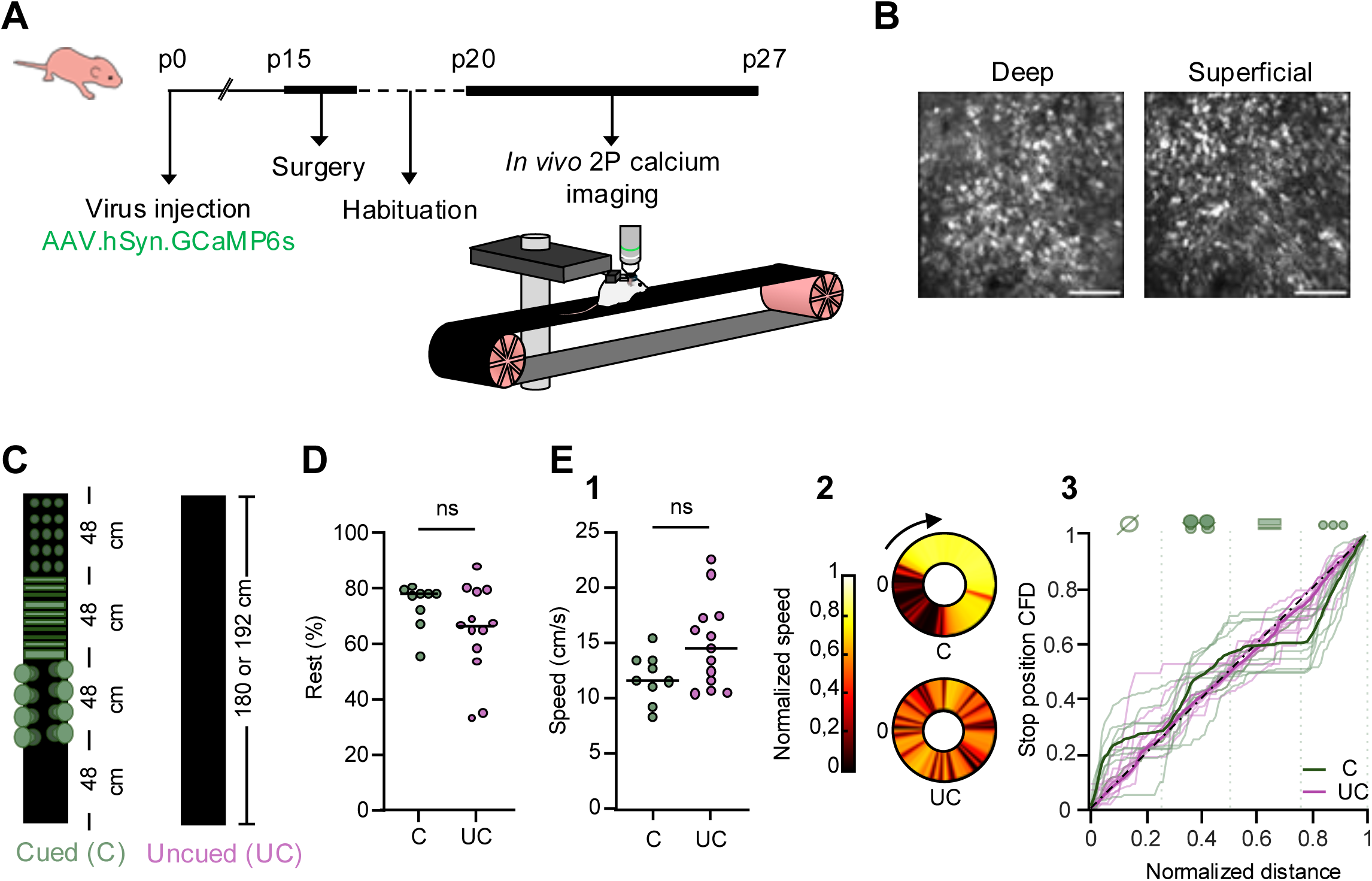
Spontaneous behavior of head-fixed mice imaged on cued (C) and uncued (UC) treadmills during the fourth postnatal week **(A)** Schematic representations of the experimental timeline and experimental setup. **(B)** Representative example fields of view (FOVs) of the GCaMP fluorescence signal from the deep and superficial CA1 *stratum pyramidale* imaged in a P23 mouse. Scale bar, 100 µm. **(C)** Schematic representation of the cued (C) and uncued (UC) treadmills. The cued treadmill is divided into four equal-sized zones: three zones with different tactile cues (from top to bottom: disks of Velcro, heat-shrink tubing, paintbrush bristles), and one cue-poor zone. **(D)** Mean time spent at rest on the cued (C, green, median = 78 % [iqr = 9], N = 9 mice) and uncued (UC, pink, median = 67 % [iqr = 32], N = 13 mice) treadmills. No significant difference was observed between the two groups (UC vs. C: Mann-Whitney test, U = 40, p = 0.2349). Each dot represents the mean rest time per animal across all recording sessions. Black lines indicate median values. **(E)** (1) Mean running speed on the cued (C, green, median = 11.6 cm/s [iqr = 3.3], N = 9 mice) and uncued (UC, pink, median = 14.5 cm/s [iqr = 6], N = 13 mice) treadmills. No significant difference was observed between the two groups (UC vs. C: Mann-Whitney test, U = 33, p = 0.0956). Each dot represents the mean speed per animal across all recording sessions. Black lines indicate median values. (2) Representative example of mean running speed distribution within session on a cued (C, top) or uncued (UC, bottom) treadmill from two different mice. Mean speed was binned and normalized to the maximum mean speed within the session. Arrow indicates the direction of treadmill rotation. (3) Cumulative frequency distribution (CFD) of stop positions across all sessions along the cued (C) and uncued (UC) treadmills. CFDs were compared to a uniform distribution (0-1, black dashed line) of matching sample size. The distribution shows preferred stop positions for the cued condition (C, green; Kolmogorov-Smirnov test: D = 0.176, p < 0.001), but not for the uncued (UC, pink; Kolmogorov-Smirnov test: D = 0.0335, p = 0.066). Thick lines represent the cumulative distributions from all animals across all sessions, and thin lines indicate individual animals.

We first characterized place-cell development as a baseline measure of externally anchored spatial representations against which to compare internally generated sequences. To monitor place-cell-like activation patterns, we first performed a series of experiments on mice placed on a treadmill enriched in salient tactile cues. The treadmill had a 192-cm belt with four zones, one deprived of local cues (cue-poor zone) and the others enriched with three different types of tactile cues (Figure 1C and methods). The influence of distal visual cues was minimized by having the mice run in the dark. Of note, mice were neither food nor water deprived, and their behavior was not guided by any external reward, as previously described^11^.

The behavioral analysis (Figure 1D-E and S1) indicates that, by the start of the 4th postnatal week, mice already display cue-guided spatial behavior, which may be supported by emerging spatial representations, as previously observed ^21–24,33^. To test this, we next searched for “place cells” from the pool of active neurons. Activity rate was significantly different across CA1 sublayers (Methods), with deep CA1 neurons displaying a higher frequency of transients than superficial ones (Linear Mixed Models (LMM), F_(1,6549.9)_ = 168.03, p < 0.0001, Table S1), consistent with previous reports in adult mice ^26^. Event rates showed small but significant differences across days (LMM, F_(7,6550.6)_ = 9.33, p <0.0001, Table S1), but the overall layer difference was already present at P20 and remained stable throughout development. We next identified place cells (Figure 2A, top panel, see Methods) and found that 24% of active neurons qualified as place cells (Figure 2B1). The proportion of “place cells” varied significantly with age (LMM, F_(3,89.8)_ = 3.74, p = 0.014), though without any particular monotonic trend (Figure 2B2, left panel) and with no difference between layers. In contrast, when computing spatial information within all active cells (see methods), we found that the superficial CA1 contained more spatial information than the deep one (LMM, F_(1,6551.2)_ = 30.53, p < 0.0001, Table S3). In order to better understand the developmental evolution of the spatial information contained in the firing of early “place cells”, we next analyzed the number, size and position of the place fields by fitting their calcium fluorescence signals (see Methods, Figure 2A, S2A). Overall, 21% of our “place cells” displayed multiple place fields. This proportion did not vary significantly with age (LMM, F_(3,90.7)_ = 2.35, p = 0.077, Figure 2B2, left panel), but was different between the superficial and deep CA1 layers, with the latter containing more multiple field place cells (F_(1,89.4)_ = 4.98, p = 0.028, Figure 2B2, right panel). In contrast, we observed a significant widening of place fields with age (LMM, F_(3, 10695)_ = 29.02, p < 0.0001; Figure 2B3), more pronounced in the superficial layer (LMM age × plane interaction: F_(3,10756)_ = 6.46, p = 0.0002; Figure S2B). Across age groups, place field distribution was uneven among treadmill zones (LMM, F_(3,43037)_ = 1030.99, p < 0.0001), with the cue-poor zone being the least covered (mean AUC = 13.4, Figure 2C-D). This distribution changed significantly with age (LMM, F_(3,42923)_ = 81.14, p < 0.0001; Figure 2D, left panel). The earliest developmental periods (P20-P21 and P22-P23) differed significantly from the latest period (P26-P27, p < 0.0001 and p < 0.0001, respectively), while the intermediate period (P24-P25) was comparable to P26-P27 (p = 1.0). Critically, we observed a significant age x zone interaction (F_(9,43037)_ = 4.56, p < 0.0001), driven notably by developmental changes in the cue-poor-zone zone. Place field representation within this zone increased substantially starting at P24-25 (Post-hoc analysis, P20-21 vs P24-25: p < 0.0001; P22-23 vs P24-25: p = 0.0005), with continued growth through P26-27 (P24-25 vs P26-27: p = 0.0470; Figure 2D, right panel). This developmental trajectory was consistent across both deep and superficial CA1 sublayers (LMM, age x zone x plane interaction: F_(9,43037)_ = 0.77, p = 0.65). Of note, the anatomical distance between CA1 “place cells” did not correlate with their place field separation (Figure S2C) with no significant effect of age (LMM, F_(3,92.53)_ = 0.105, p = 0.957) or sublayers (LMM, age x plane interaction: F_(3,89.17)_ = 0.771, p = 0.514). We conclude that early place cells rely on salient cues until P24 and progressively widen their place fields.

**Figure 2.**
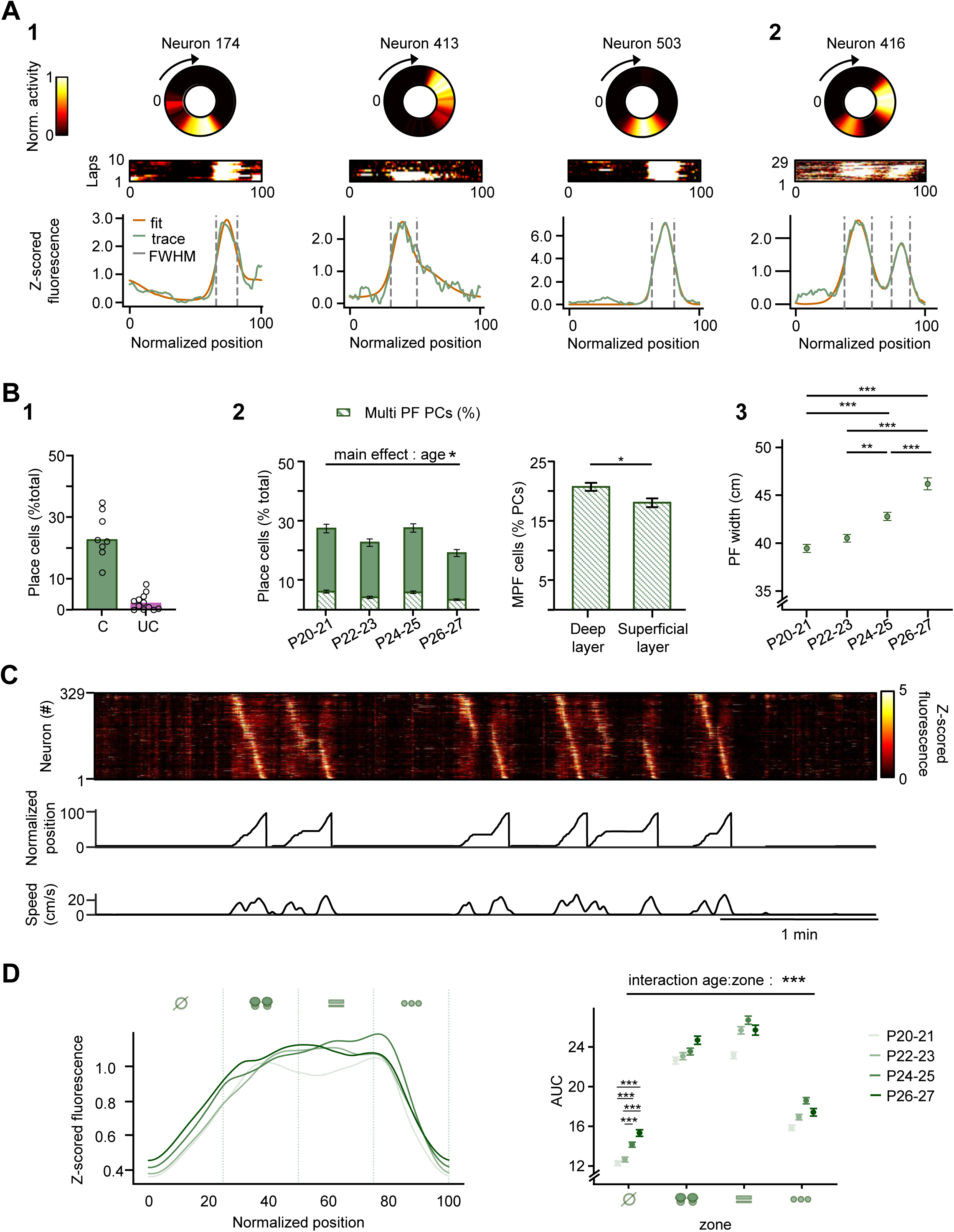
Developmental evolution of cue-based CA1 representations at single-neuron level **(A)** Illustration of CA1 place cells and their spatial tuning via Von Mises (VM) mixture fits. (1) Top: three representative examples of place cells from the same recording session. Circular heatmaps display normalized mean fluorescence activity of individual neurons as a function of normalized mouse position on the track. Each circular map is paired with its corresponding lap-by-lap rate map. Bottom: Z-scored fluorescence traces (green) of the same neurons plotted against normalized position, fitted with a von Mises distribution (orange; see Methods) used as a proxy for the place field. The grey line indicates the full width at half maximum (FWHM), which serves as a proxy for place field width. (2) Same as (1) but for a representative neuron from a different recording session exhibiting multiple place fields. **(B)**(1) Percentage of place cells normalized by the number of active cells in the field of view (FOV) for cued (green, C, median = 23 % [iqr=12.5], N = 8 mice) and uncued (pink, UC, median = 2 % [iqr = 4], N = 12 mice) conditions. Each data point represents an individual animal. (2-3) Analyses in the cued condition only. (2) Left panel: Place cell percentages across age groups (same normalization as in (1)). Data are presented as mean ± SEM (P20-21: 27 ± 1.9%, n = 22 sessions, N = 7 mice; P22-23: 23 ± 1.8%, n = 32 sessions, N = 8 mice; P24-25: 28 ± 2.0%, n = 26 sessions, N = 7 mice; P26-27: 20 ± 1.8%, n = 24 sessions, N = 7 mice; See Table S2 for detailed statistics). The hatched portion of each bar represents the percentage of place cells exhibiting multiple place fields (P20-21: 6 ± 0.6%; P22-23: 4 ± 0.5%; P24-25: 6 ± 0.6%; P26-27: 4 ± 0.4% (same n and N as above; See Table S4 for detailed statistics). Right panel: Percentage of place cells with multiple fields among all place cells in the deep (21 ± 0.9%) and superficial layers (18 ± 1%; n = 52 sessions N = 8 mice for both, See Table S4 for detailed statistics). (3) Place field width across age groups (P20-21: 39 ± 0.4cm n = 2923 PCs N = 7 mice, P22-23: 41 ± 0.4cm n = 3106 PCs N = 8 mice, P24-25: 43 ± 0.4cm n = 2889 PCs N = 7 mice, P26-27: 46 ± 0.6cm n = 1851 PCs N = 7 mice; See Table S5 for detailed statistics). **(C)** Example of population activity of CA1 place cells (deep and superficial combined) recorded on the cued treadmill. Each row shows the z-scored calcium activity of a single neuron across several laps, plotted over time. Cells are sorted according to the position of their peak activity in the mean activity across laps. The animal’s normalized position (middle) and running speed (bottom) are shown for the same time window. **(D)** Left panel: Mean place field fits (as shown in Figure 2A1, bottom panel, orange) for each age group, plotted along normalized position in the cued condition. Position is divided into four segments corresponding to the cue belt configuration (from the left to the right: cue-poor zone (z1), paintbrush bristles (z2), heat-shrink tubing (z3), disks of Velcro (z4). Right panel: Mean area under the curve (AUC) for each zone across age groups (z1: P20-21: 12 ± 0.2; P22-23: 13 ± 0.3; P24-25: 14 ± 0.3; P26-27: 15 ± 0.3. z2: P20-21: 23 ± 0.3; P22-23: 23 ± 0.3; P24-25: 24 ± 0.3; P26-27: 25 ± 0.4. z3: P20-21: 23 ± 0.3; P22-23: 26 ± 0.4; P24-25: 27 ± 0.4; P26-27: 26 ± 0.5. z4: P20-21: 16 ± 0.3; P22-23: 17 ± 0.3; P24-25: 19 ± 0.3; P26-27: 17 ± 0.4. For all age groups: n = 2923, 3106, 2889, and 1851 PCs from N = 7, 8, 7, and 7 mice, respectively; see Table S6 for detailed statistics). Statistical comparison performed using Linear mixed model (random effect: mouse identity); *p < 0.05, **p < 0.01, ***p < 0.001.

Finally, we probed the stability of position representation at single-cell and population levels using Track2p^29^, a novel algorithm specifically designed for longitudinal tracking of neural activity across several days during early postnatal development (Figure 3A). In total, we tracked 215 place cells across 6-8 days (N = 6 mice). The place cell population displayed a significant turnover, with on average, only 13.4 ± 2.7% of the neurons qualifying as “place cells” across consecutive sessions, with no developmental trend. The correlations between place field positions for neurons identified as “place cells” on at least two days varied with age (LMM, F_(27,1784.5)_ = 2.6, p < 0.001). Pearson correlations increased progressively from early to late postnatal periods, with P26-P27 showing significantly higher stability than P20-P21 (p = 0.043; Figure 3B, middle panel). Remarkably, neurons classified as place cells at both P20 and P24 showed complete remapping (r = 0.06), whereas place fields remained quite stable between P24 and P27 (r = 0.76), with these correlations differing significantly (p = 0.0006). Overall, early (P20 to P24) and late (P24 to P27) developmental periods differed significantly (LMM, F_(1,1150.5)_ = 5.33, p = 0.02; Figure 3B, right panel), indicating the mid-fourth postnatal week as a critical transition age with substantial spatial reorganization across both layers (age period × sublayer interaction, F_(1,1146.7)_ = 0.13, p = 0.7; Figure S3A).

**Figure 3.**
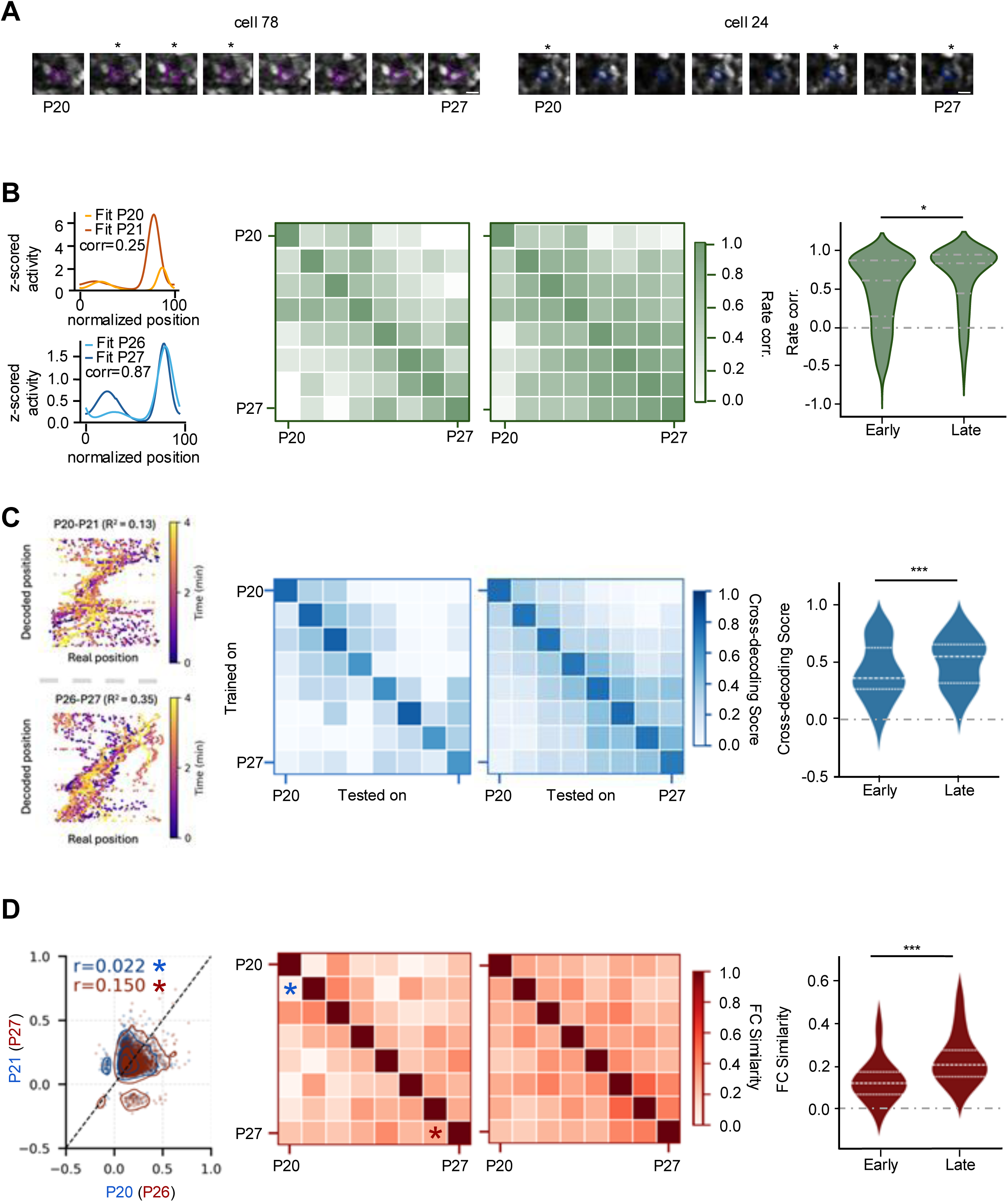
Longitudinal tracking and stability of CA1 spatial and functional representations in mice under cued conditions **(A)** Examples of cells tracked throughout the entire recording period (P20-P27) and identified as place cells (*) on three consecutive days (left) and on three intermittent days (right). Scale bar, 10 µm. **(B)** Top left: example of place fields correlation for a single representative cell for an illustrative animal between days P20 (yellow) vs. P21 (orange). Bottom left: same as top left, shown for a single representative cell between days P26 (light blue) vs. P27 (dark blue). Middle left: Pearson correlation for all combinations of pairs of days in the example animal. Middle right: Mean pearson correlation for all day pairs averaged across all animals (N = 6 mice, see Table S8 for detailed statistics). Right: Distribution of all cross-day correlation values for early pairs of days (between P20 and P23, median = 0.6, [iqr = 0.7], n = 763 observations, N = 6 mice) compared to late pairs of days (P24-P27, median = 0.8, [iqr] = 0.5, n = 392 observations, N = 6 mice). See Table S9 for detailed statistics. **(C)** Left: example cross day position decoding for the same animal as in (B) for a pair of early days (P20-P21) and a pair of late days (P26-P27). Coefficient of determination (R^2^) values are indicated. Middle left: R^2^ values for all combinations of days used in cross-day decoding in the example animal. Middle Right: Mean R^2^ values for all combinations of days averaged across all animals (N = 6 mice). Right: Distribution of all cross-day decoding values for early days (between P20 and P23, median = 0.35, [iqr = 0.38], n = 77 observations, N = 6 mice) compared to late days (P24-P27, median = 0.54, [iqr = 0.36], n = 82, N = 6 mice, see Table S10 for detailed statistics). **(D)** Left: Example of functional connectivity (FC) comparisons for the same animal as in (B) and (C) between days P20 vs. P21 (blue) and P26 vs. P27 (red). Spearman correlation coefficients (FC Similarity, FCS) are indicated. Middle left: FCS values for all day pairs in animal 17. Middle right: Average FCS across all mice. Right panel: Distribution of all unique FCS values (one per day pair, lower triangle only) for early (P20-P23, median = 0.12, [iqr = 0.12], n = 30 observations, N = 6 mice) and late (P24-P27, median = 0.20, [iqr = 0.13], n = 33 observations, N = 6 mice) periods (see Table S11 for detailed statistics). Statistical comparison performed using Linear mixed model (random effect: mouse identity); *p < 0.05, **p < 0.01, ***p < 0.001.

After quantifying spatial coding properties and their stability at the level of single cells, we turned to population level given that neurons not classified as place cells can also include spatial information^34^. We first performed PCA embeddings of neural activity during run epochs, which showed a ring-like topology across all days, recapitulating the circular topology of the treadmill (Figure S3C2). This correspondence between position and activity was also made clear when calculating the spatially averaged fluorescence of all neurons and sorting cells based on their maximum fluorescence (see Methods, Figure S3C1). When projecting activity to PCA space across days we saw a significant loss of topology, suggesting an unstable relationship between neural activity and position (Figure S3C2). A similar breakdown of structure was observed when re-sorting the spatially averaged fluorescence of a given day based on the sorting obtained on a different one (Figure S3C1). To quantify these effects we turned to regression analysis (see methods), to decode the actual position of the mouse from the simultaneously recorded population activity (Figure 3C, left panel and Figure S3C3, left panel). Interestingly, despite changes to single cell coding properties, the within-day decoding performance was stable across development (LMM, p<0.0001, Figure S3B). Tracking the same neurons across days, we could also probe the stability of this representation by fitting a model on one day and testing it on all other days (Figure 3C, middle panel; Figure S3C3). Interestingly, this analysis indicated a higher performance of cross-day decoding for later developmental epochs (LMM, F_(1,155.35)_ = 13.459, p < 0.0001, Figure 3C, right panel), suggesting a developmental stabilisation of population coding properties, consistent with our results on the stabilisation of place field properties of single neurons.

Our findings demonstrate that cue-based position coding is present in both deep and superficial CA1 pyramidal layers by the beginning of the fourth postnatal week in mice. However, this spatial representation remains unstable and dependent on salient extrinsic cues until postnatal day 24 (P24). The functional maturation of CA1 position coding observed at P24 prompted us to investigate whether this developmental milestone coincides with an overall stabilization of the functional structure of the CA1 network. To this aim, we analyzed the cross day similarity in the functional connectivity between active CA1 neurons, quantified as the Pearson correlation across all neuron pairs, after filtering for statistical significant connections only (the resulting adjacency matrices had densities in the range 0.50 ± 0.25; See Methods). Similarity values were significantly higher in the late period (P24-27) compared to the early period (P20-23, LMM, F_(1,57.69)_ = 18.26, p < 0.0001, Figure 3D). This result points to a progressive slowing down of the network reorganisation towards the end of the fourth postnatal week, with a significant shift occurring from P24 onwards.

This, together with the protracted emergence of “place cells” in the cue-poor portions of the treadmill led us to investigate the emergence of self-referenced internal sequences. Such internal sequences could potentially bridge spatial gaps between discrete cues and represent a critical indicator of maturing CA1 internal dynamics, reflecting the transition from externally-driven to internally-generated hippocampal representations.

### Protracted development of internal sequences

To address this question, we performed similar imaging experiments in mice alternating between rest and running at various speeds on a belt devoid of any cues (Figure 1). As quantified above, mice did not display any preference for a specific location on the belt in these conditions (Figure 1E3) and position-selective activity of CA1 neurons was exceptionally rare (median percentage of place cells: 1.85%, [iqr = 3.92], N = 12 mice, Figure 2B1). Previous work, in the same conditions but in adult mice, had shown that CA1 dynamics organize into recurring sequences of neuronal activation integrating self-referenced information in the form of run distance or elapsed time ^10,11^. We used our previously established method to detect recurring sequences during run periods (^11^, see Methods) and found that these internally-recurring sequences followed a delayed developmental trajectory compared to cue-driven CA1 firing, with a distinct transition at P24 (Figure 4A). A developmental increase in sequence detection was observed across imaging sessions. Using generalized linear mixed models (GLMM), we found a significant effect of developmental period on the likelihood of detecting sequences (Wald χ² = 6.87, df = 2, p = 0.032). Sequences were detected in only 6% of imaging sessions during the early period (P20-P23) compared to 26% during the later period (P24-P27; z = −2.28, p = 0.034, Figure 4A4). This developmental increase was independent of the imaging plane, occurring similarly in both superficial and deep layers (plane effect: Wald χ² = 0.35, p = 0.555; period × plane interaction: Wald χ² = 1.25, p = 0.534).

**Figure 4.**
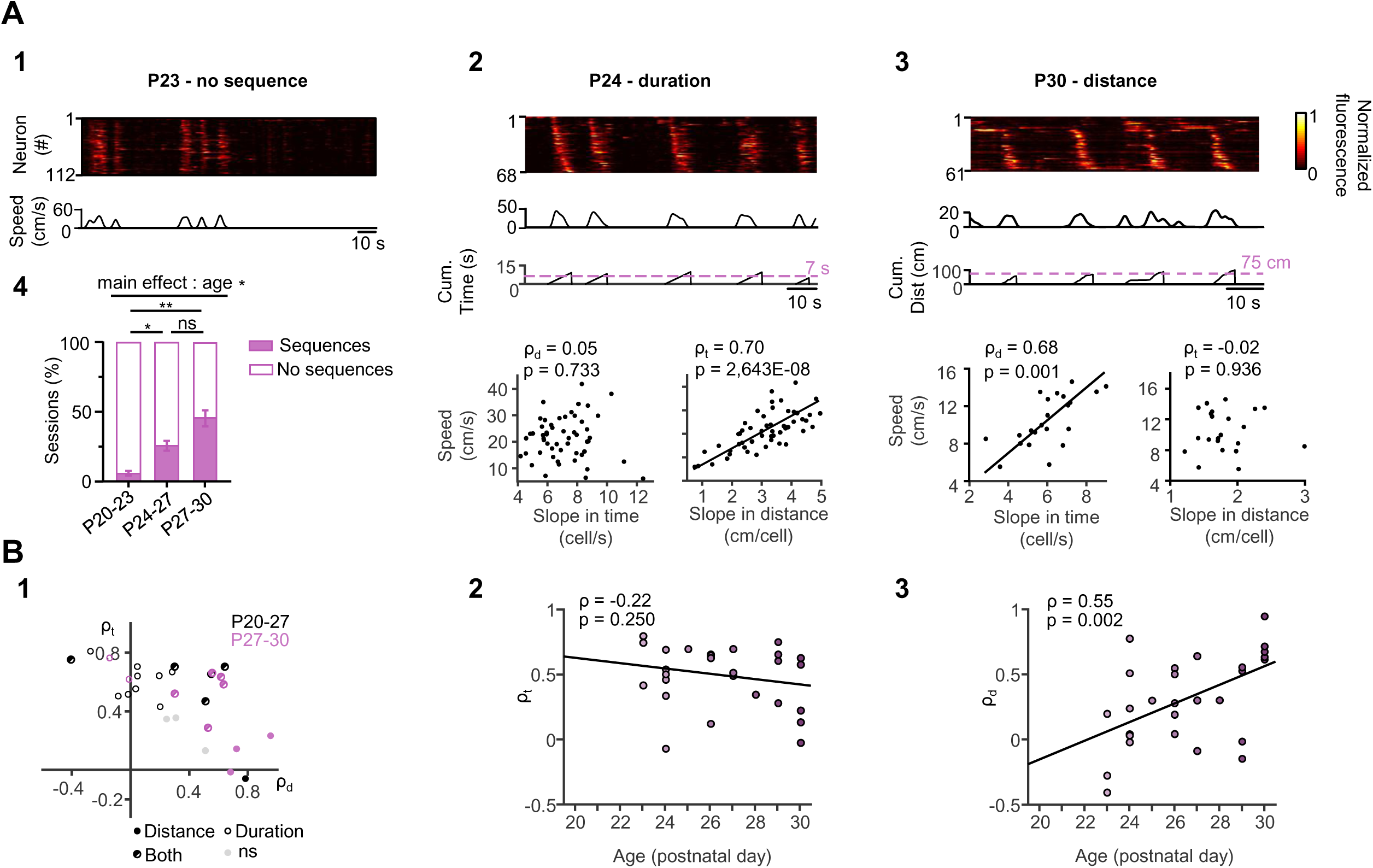
Emergence of internal sequences during the fourth postnatal week **(A)**(1) Example of calcium activity recorded from a P23 mouse running on the uncued treadmill. Rasterplot showing the normalized fluorescence activity of neurons active during the running period as a function of time (top). Running epochs are defined by speeds above 2 cm/s (bottom). (2) Same as (1) for a P24 mouse. The raster plot shows neurons organized in sequences during running epochs (top). The cumulative duration of these sequences indicates temporal consistency across sequences (middle). Relationship between speed and temporal slope for each detected sequence (dots) recorded in the same imaging session (bottom left). Relationship between speed and spatial slope for the same sequences (bottom right). On each plot, the black line indicates the linear fit with the associated Spearman correlation coefficient and p value. (3) Same as (2) for a P30 mouse. The cumulative distance of these sequences indicates consistency across distance (middle). (4) Proportion of session displaying internal sequences (pink) at P20-23 (6 ± 3% n = 49 sessions, N = 11 mice), P24-27 (26 ± 6 %, n = 54 sessions, N = 5 mice) and P27-P30 (46 ± 10 %, n = 24 sessions, N = 3 mice). Data are presented as mean ± SE and statistical comparison performed using Generalized Linear Mixed Model (random effect: mouse identity); *p < 0.05, **p < 0.01. See Table S12 for detailed statistics **(B)**(1) Across all imaging sessions (P20-27: n = 17 sessions, N = 5 mice; P27-30: n = 11 sessions, N = 3 mice), Spearman correlation coefficients for time (ρₜ) are plotted as a function of Spearman correlation coefficients for distance (ρ_d_). Sessions recorded between P20-27 and P27-30 are represented by black and pink dots, respectively. Sessions showing significant correlations for distance, time, both, or neither are indicated by filled, open, half-filled, or gray dots, respectively (2) Spearman correlation for time (ρ_t_) as a function of age (P20-P30). Each dot represents an imaging session, and the black line shows a linear fit with Spearman correlation coefficient and p value associated. (3) Same as (2) for Spearman correlation for distance (ρ_d_).

Internally-recurring sequences engaged a sparse and scattered neuronal population representing a median of 29% of active neurons at P20-P23 and 21% at P24-P27, with no significant developmental difference (Mann-Whitney test, p = 0.20; Figure S4A). Similarly, sequence frequency during run periods did not differ between age groups (P20-P23: median = 0.06 Hz; P24-P27: 0.08 Hz; p = 0.35; Figure S4A). Cells participating in these sequences were spatially intermingled with other active cells displaying variable activity patterns (not shown).

Although these results indicate that internally-recurring sequences develop later than cue-driven representations, this delayed emergence could reflect the impact of experience rather than a developmental process per se. To distinguish between these alternatives, we imaged mice starting at P27 (n = 3 mice). These mice underwent the same treadmill habituation as younger animals prior to their first imaging session. Sequences were observed in 46% of imaging sessions over P27-P30, comparable to P24-P27 (z = −1.44, p = 0.150; Figure 4A4), indicating that the delayed development of internal sequences reflects age-dependent maturation rather than progressive habituation to the treadmill.

We next asked whether these internally generated sequences could encode run duration or distance. To do so, we computed, for each imaging session with sequences, the correlation between the slopes of the sequences (in the temporal or spatial domains), and the median mouse speed of the corresponding run epochs. If the progress of neuronal activation was set by the run distance, then the temporal slope of the sequences (expressed as cell/s) should correlate with speed while the spatial slope (in cm/cell) should be constant and thus independent from speed. We calculated, for each imaging session, the Spearman correlation coefficient between the temporal or spatial slope and the speed (Figure 4A2 and 3, bottom panels). Between P20 and P27, most sequences encoded elapsed time (9/17 sessions with sequences, Figure 4B) rather than distance (1/17 sessions with sequences, Figure 4B). Moreover, a mixed spatiotemporal representation ^10^ was already present between P20 and P27 (5 of 17 sessions with sequences, Figure 4B1). Interestingly, the proportions of duration-, distance-, and mixed- coding sequences changed over development. At P27-30, most sequences displayed a mixed representation of time and distance (5 of 11 sessions with sequences) with two sessions displaying duration-coding and three distance-coding sequences. This developmental evolution was mirrored by a significant increase in distance coding with age (Figure 4B3), while duration coding remained stable (Figure 4B2). Comparable results were obtained when considering only significant sequences per run epoch instead of sessions (see Methods, Figure S4B).

Altogether, these results demonstrate that the internal organization of CA1 activity develops gradually from P24, independently of experience. The information supported by these internally recurring sequences continues to evolve beyond the fourth postnatal week, progressively acquiring adult-like properties.

### Cell assemblies develop before internal sequences with layer-specific expansion patterns

We previously demonstrated that recurring internally-generated sequences in adult mice comprise cell assemblies observed during awake rest periods^13^. However, the directionality of this relationship remains unclear: do pre-existing cell assemblies provide a functional scaffold for the development of sequences, or do sequences emerge first and their segmentation drive the formation or reactivation of cell assemblies? To address this question, we examined the developmental emergence of cell assemblies focusing on rest episodes (Figure 5). In both cued and uncued conditions, we found that CA1 dynamics during rest periods are organized into synchronous calcium events, involving a proportion of active cells that exceed chance levels (SCEs, see Methods; Figure 5A-B). SCEs occurred in both cued and uncued conditions, in superficial and deep CA1, as early as P20. SCE frequency did not significantly differ between P20-23 and P24-27 (not shown, LMM, F_(1,129.1)_ = 0.68, p = 0.41, Table S13), regardless of the condition (cued or uncued) or CA1 sublayer. In contrast, SCE amplitude (percentage of active cells involved in SCEs) showed an increase with age (LMM, F_(1,130.9)_ = 4.23, p = 0.04; not shown, Table S14). Still no significant interactions with condition or sublayer were detected, indicating that age-related changes in SCE amplitude were not specific to either cued or uncued conditions, nor restricted to particular layers.

**Figure 5.**
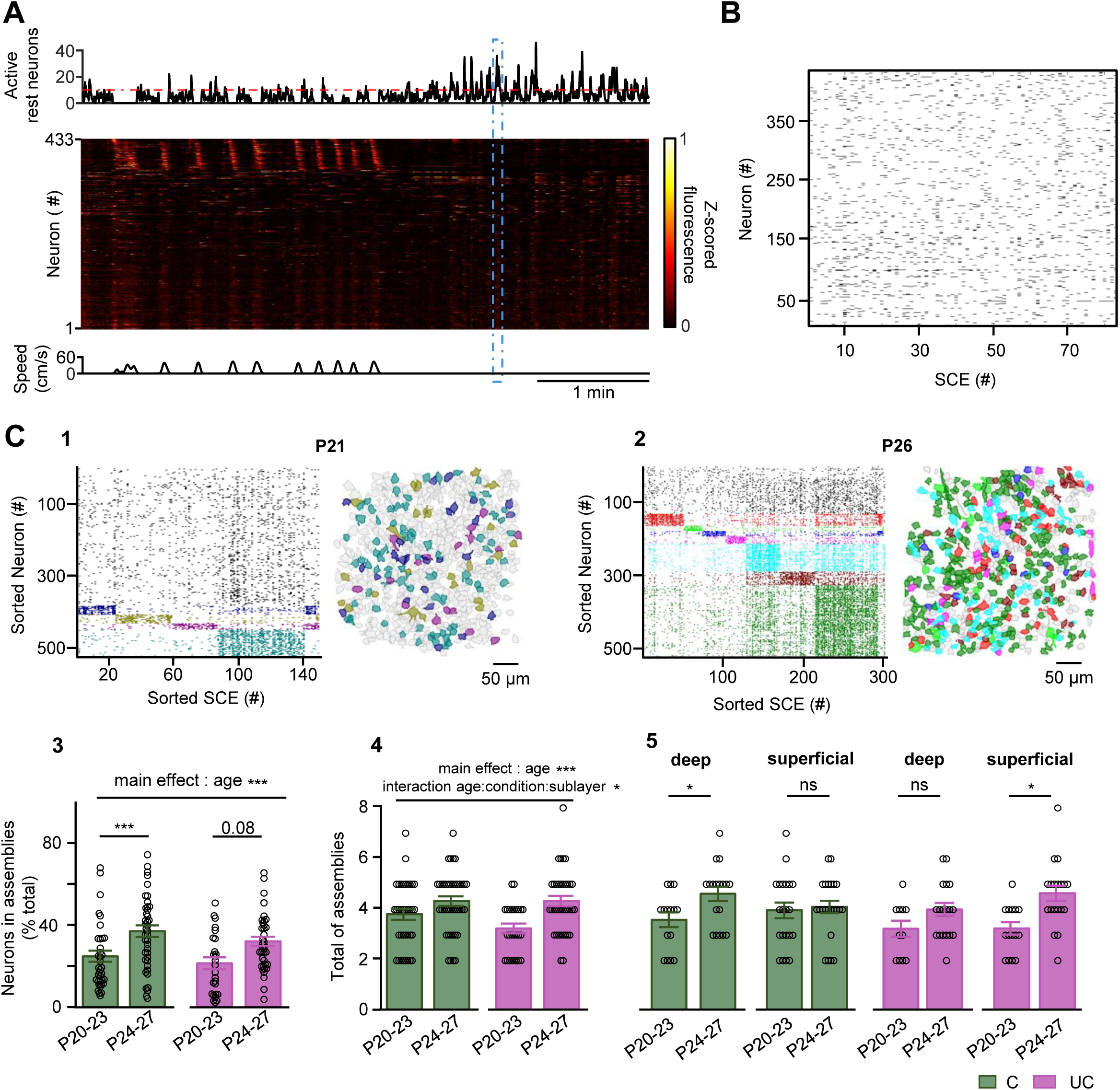
Detection and characterization of synchronous calcium events and neuronal assemblies **(A)** Detection of a synchronous calcium event (SCE) during a rest period. Top panel: Number of active neurons per frame during rest periods over a given time period. The red dashed line represents the detection threshold set at 10 active neurons (see Methods). Middle panel: Heat map of neuronal calcium activity over time. Each row shows the z-scored calcium activity normalized of a single neuron across several laps, plotted over time. Bottom panel: speed of the mouse (cm/s) over time. Blue rectangle: Delimits a detected SCE confirmed across all three panels. **(B)** Raster plot of neuronal participation in detected SCEs. Each row represents an individual neuron, and the x-axis shows the detected SCEs. **(C)**(1) Clustering of neurons into functional assemblies based on SCE activity in a single mouse at P21. Left: Raster plot of neuronal activity during SCEs, sorted and color-coded by cluster membership. Each color represents a distinct cluster. Right: Spatial localization of neurons in the field of view (FOV), with each neuron colored according to its cluster assignment (scale bar: 50 µm). (2) Same as (C)(1) for the same mouse at P26. (3) Percentage of neurons engaged in assemblies (normalized by total active neurons in the FOV) for early (P20-P23) and late (P24-P27) periods. Data are presented as mean ± SE and are shown for cued (C, green; early: 25 ± 2.7 % n = 36 sessions N = 7 mice; late: 37 ± 2.8 % n = 41 sessions N = 8 mice) and uncued (UC, pink; early: 21 ± 2.9 % n = 26 sessions N = 8 mice; late: 32 ± 2.3 % n = 37 sessions N = 6 mice; See Table S16 for detailed statistics) groups. Each point represents a single recording session. (4) Number of detected assemblies for early (C: 3.8 ± 0.2 assemblies; UC: 3.2 ± 0.2 assemblies) and late (C: 4.3 ± 0.2 assemblies; UC: 4.3 ± 0.2 assemblies) periods (same n and N as above; See Table S17 for detailed statistics). Each point represents a single recording session. (5) Same as (4) for deep ( C, green: early: 3.5 ± 0.3 assemblies n = 15 sessions N = 6 mice, late: 4.5 ± 0.3 assemblies n = 18 sessions N = 7 mice; UC, pink: early: 3.2 ± 0.3 assemblies n = 11 sessions N = 6 mice, late: 3.9 ± 0.3 assemblies n = 18 sessions N = 6 mice) and superficial ( C, green: early: 3.9 ± 0.3 assemblies n = 21 sessions N = 7 mice, late: 4 ± 0.2 assemblies n = 23 sessions N = 8 mice; UC, pink: early: 3.2 ± 0.2 assemblies n = 15 sessions N = 8 mice, late: 4.6 ± 0.3 assemblies n = 19 sessions N = 5 mice; See Table S17 for detailed statistics) layers. Statistical comparison performed using Linear mixed model (random effect: mouse identity); *p < 0.05, **p < 0.01, ***p < 0.001.

We next detected cell assemblies nested in SCEs using a k-means-based method (Figure 5C), as previously reported^13^. Cell assembly organization was present throughout the developmental period studied here, even when internal sequences were absent. The proportion of sessions exhibiting at least two cell assemblies remained stable across ages (P20-23 vs. P24-27), conditions (cued vs. uncued), and CA1 layers (deep vs. superficial). A generalized linear mixed model (GLMM) revealed no significant main effects of age (χ²(1) = 0.17, p = 0.68), condition (χ²(1) = 0.52, p = 0.47), or CA1 plane (χ²(1) = 3.72, *p*= 0.054), and no significant interactions (all *p* > 0.47, see Table S15). Overall, cell assemblies were detected in more than half of the sessions, suggesting that the local circuits shaping synchronous population events into structured assemblies are already wired by P20. Remarkably, the percentage of neurons involved in assemblies significantly increased with age (LMM, F_(1,129.7)_ = 16.83, p < 0.001; Figure 5C3). Post-hoc comparisons revealed that this age-related increase was significant in cued conditions (p < 0.001) but only a trend in uncued conditions (p = 0.08). No significant layer-specific differences or interactions were observed (Figure 5C3). The absolute number of assemblies detected per session also significantly increased with age (LMM, F_(1,129.3)_ = 12.03, p < 0.001), with a significant age x sublayer x condition interaction (F_(1,119.5)_ = 4.05, p = 0.046; Figure 5C4). Post-hoc comparisons revealed that this developmental increase in assembly number was specific to cued conditions in the deep layer (p = 0.032) and uncued conditions in the superficial layer (p = 0.038), while no significant changes were observed in the other sublayer-condition combinations (Figure 5C5). We also examined the anatomical organization of cell assembly members and found that overall cells engaged in assemblies were spatially intermingled with other cells. Linear mixed models revealed no significant variation in this spatial intermingling pattern across age groups, layers, or experimental conditions (LMM, all p > 0.05; Table S18).

We then analyzed the recruitment of place cells and internal sequence neurons into cell assemblies. Linear mixed models revealed a significant effect of belt type on the proportion of neurons recruited in assemblies (LMM, F_(1,33.166)_ = 7.00, p = 0.012; Table S19), with a higher recruitment of place cells than neurons involved in run sequences in rest assemblies. When examining the reverse relationship (assembly neurons involved in sequences or place cells), a similar trend was observed (LMM, F_(1,16.406)_ = 3.41, p = 0.083; Table S20). No significant effect of age, CA1 layer, or their interactions was detected across any of these analyses, indicating that this relation between neuronal involvement in run sequences and rest assemblies was stable across development and consistent across sublayers. Since assemblies are observed prior to sequences in uncued conditions, we last tested if sequences could emerge from previously co-active neurons in uncued conditions. To this aim, we tested whether cell pairs recruited in internal sequences on a given day (day 1) were more correlated during rest periods recorded the day before (day 0). For this, we restricted our analysis to cases where sequences could be found on day 1 and assemblies on day 0 in the same layer and mouse (n=10 day pairs). If in 40% of the cases, we could observe significantly more correlated rest activity on day 0 in the pairs of cells that will be involved in sequences on day 1 (p-values ranging from < 0.001 to 0.025, not shown), in the majority of the cases there was no significant difference in rest correlation (p-values ranging from 0.15 to 1.0). We found that internally-recurring sequences can emerge from previously co-active cell pairs, but this mechanism does not seem to represent a general developmental trend.

## Discussion

Using longitudinal calcium imaging in head-fixed mice, we tracked the day-by-day emergence of CA1 sequences and cell assemblies during the third postnatal week, a critical period that marks the onset of experience-dependent plasticity in hippocampal circuits. Our findings reveal a discrete developmental transition at postnatal day 24, characterized by the stabilization of cue-based position coding and functional connectivity, the expansion of cell assemblies and the emergence of internally-recurring sequences. While cue-based spatial representations are present from the earliest time points examined, these P24 transitions should collectively contribute to the stabilization of the CA1 cognitive map, which continues evolving after the 4th postnatal week with the progressive increase in distance coding.

We have also uncovered emergent laminar organization in the developing CA1. Deep neurons are more active, more likely to exhibit multiple place fields and display less spatial information than their superficial counterparts. The developmental evolution of rest assemblies is also sublayer specific, as their number increases only in deep CA1 in the cued treadmills and in superficial CA1 in the bare ones. These layer-specific patterns suggest the early emergence of radial functional differentiation, but the overall developmental trajectory (including the P24 transition) remains largely shared across layers. Together, these findings provide a comprehensive multidimensional description of CA1 circuit maturation and establish the third postnatal week as a critical vulnerability window for hippocampal cognitive map development.

### Mechanistic and functional significance of the mid-fourth postnatal week transition

We provide the first evidence that externally anchored representations emerge before internally driven, self-centred dynamics such as duration and/or distance-coding sequences. We also show that cell assemblies observed during rest develop prior to internally-recurring run sequences. This ordering suggests that stable external reference frames scaffold the subsequent development of internally generated activity. While this appears at odds with the fact that allocentric navigation emerges later than egocentric strategies ^27^, it is consistent with the finding that children can use landmark-based strategies but cannot yet integrate landmark and self-motion information ^28^. Thus, internally-generated hippocampal dynamics do not reflect a more basic version of cue-based representations. Instead, they may reflect higher-order cognitive computations such as integration of egocentric and allocentric information or relational mapping beyond spatial variables ^4,35^. Hence, the fourth postnatal week marks not only maturation of spatial coding but also the emergence of internally generated dynamics that may underpin flexible spatial behavior and more precise episodic-like memory ^36^.

These functional changes at P24 are supported by maturation of both local and long-range circuits. Within CA1, parvalbumin-positive basket cells and their perineuronal nets refine substantially ^36,37^, and GABAergic synapses between interneurons acquire activity-dependent potentiation only after P24 ^38^. This inhibitory maturation enables selective recruitment of pyramidal cells during sharp-wave ripples and provides the perisomatic drive necessary for organizing sequences that encode behavioral trajectories ^23^. Competitive allocation mediated by PV-expressing interneurons further establishes sparse, precise engrams supporting episodic-like memory ^36^. Another turning point may be excitatory and inhibitory synapses becoming plastic from P23 onwards ^39^. The emergence of internal sequences may also depend on the maturation of multiple circuit elements, including superficial CA1 neurons—the latest-born pyramidal population—whose output to the entorhinal cortex is essential for grid cell function and self-referenced spatial coding, particularly in cue-poor environments ^40,41^. This delayed maturation likely contributes to the lag of internally generated sequences relative to cue-driven activity, which relies on early spontaneous network patterns (eSPWs, SCEs) and sensorimotor inputs ^17–19,42^ Beyond local circuits, inputs from CA3 and entorhinal cortex become functionally mature around P24 ^43^.

The circuit mechanisms supporting the seconds-long Internal sequences observed here should be similar to those supporting the generation of theta sequences, which emerge after P23-24 and require medial septal inputs ^9,23,25^. Awake rest replay similarly transitions around this stage from persistent firing at the current location to reactivation of progressively longer past trajectories ^23^, reflecting the coordinated maturation of septal cholinergic and GABAergic projections. The temporal alignment of theta sequence maturation, internal sequence emergence, and CA1-entorhinal loop refinement suggests that self-referenced coding depends on both intrahippocampal and extrahippocampal circuits. Interestingly, our modeling work ^10^ suggests that the shift in internally-recurring sequences from encoding duration to distance could simply result from an increased extrinsic excitatory drive (including neuromodulation) integrating speed information, without the need of further local rewiring. Together, these local, network, and systems-level changes converge at P24, transforming CA1 from a circuit supporting cue-dependent navigation into one capable of flexible, context-sensitive spatial and relational representations.

### Developmental trajectories of CA1 spatial representations: comparison with prior studies

Our findings build upon prior work examining CA1 spatial map development in rats, mostly through electrophysiological approaches ^21–23,25,33,37^, and more recently using imaging^24^. Calcium imaging offers two major advantages in this context: (1) the ability to track the same individual neurons across multiple days, and (2) precise access to both deep and superficial sublayers of CA1 pyramidal cells. When combined with our unique behavioural paradigm that reveals internally generated sequences ^11^, this approach enables simultaneous dissection of externally anchored and internally driven network dynamics, and their stabilization across time and layers, an analytical depth not achievable with previous developmental studies. In a recent imaging study ^24^, cell tracking was limited to pairs of days within early development (up to P22 in rats) due to the rapid postnatal growth of the brain. Using our longitudinal imaging approach in head-fixed mice ^29^, we overcame these constraints and tracked place cell properties across the entire fourth postnatal week.

Regarding the proportion of place cells during development, our findings reveal that while age accounts for some variability, there is no significant monotonic increase or decrease across the ages examined. This result aligns with previous electrophysiological data in developing rats ^21^ showing relatively stable place cell proportions (∼50%) between P19-P27, but contrasts with others reporting a more pronounced increase from ∼60% to 80% between P20 and P28 ^22^. We attribute these discrepancies partially to methodological differences: electrophysiological recordings sample from more restricted regions and rely on identification of active cells during sharp-wave ripples, whereas our imaging approach provides denser local sampling. Additionally, our experimental design employed unmotivated animals without food or water deprivation, which may affect place cell recruitment compared to goal-directed behavior paradigms used in some prior studies. In any case, the proportion of place cells observed here is similar to previous reports using imaging in adult rodents (from 20 −50%), indicating a rather mature landmark-based CA1 representation ^30,44,45^. Our findings are consistent with previous electrophysiological reports showing that the fourth postnatal week marks a key transition in spatial coding and sequence maturation in rats ^23,24^. This cross-species consistency supports the validity of our head-fixed paradigm and indicates that our results capture fundamental principles of hippocampal circuit maturation In contrast to ^24^, we did not observe any significant developmental bias towards an increased recruitment of deep layer early place cells in cell assemblies or SCEs compared to superficial ones. This difference may stem from different imaging conditions or the prior use of a simpler statistical model (two-way ANOVA as opposed to the LMM used here) that may not have accounted for the non-independence of repeated measurements from the same animals. Still, our results confirm that the functional subdivision of CA1 along the radial axis is overall substantially milder during early development than in mature animals (see also below). Our data also aligns with several key findings from prior electrophysiological work: the increase in the fraction of neurons recruited in SCEs and assemblies in the cue-rich conditions parallels the previously reported increase in ripple power between P22 and P27 ^23,37^, while the expansion of place field size with age also replicates previous observations ^33^.

### Emergent Radial Laminar Organization in Developing CA1

In adult CA1, a clear functional segregation exists along the radial axis of the principal cell layer, encompassing multiple metrics including activity rates, place cell prevalence and stability, sharp wave-ripple and assembly recruitment, and inhibitory drive ^8,15,16,30–32,46,47^. Strikingly, our study reveals a milder functional subdivision during development. Place cell proportions and stability, representations of cue-poor treadmill segments, and recruitment into distance-coding sequences showed no significant dependence on CA1 sublayer, even at the end of the fourth postnatal week. Nevertheless, our analysis reveals several layer-specific features that hint at emerging functional differentiation along the radial axis. First, neurons in deep CA1 are more active, most often display multiple place fields, and overall carry less spatial information (irrespective of the mouse age), in agreement with the tendency of earlier born neurons to generalize ^16,30,46,48,49^. Second, place field width undergoes more pronounced age-dependent expansion in superficial CA1, potentially reflecting ongoing refinement of spatial selectivity in the later-born neuronal population. Third, cell assembly dynamics display divergent developmental trajectories: both the number of cell assemblies and the fraction of neurons involved in them increase significantly in deep CA1 in cued conditions, whereas superficial CA1 shows corresponding increases only in uncued treadmills. This dissociation suggests that deep and superficial layers may differentially process landmark-based versus self-motion information during early development.

Hence, the full expression of laminar specialization requires not only the intrinsic maturation of pyramidal neurons born at different developmental timepoints, but also the coordinated refinement of layer-specific inhibitory microcircuits that sculpt their activity patterns and network integration.

## Conclusion

This comprehensive longitudinal analysis of CA1 circuit functional maturation reveals that hippocampal development is not gradual but rather punctuated by critical transitions that align with the emergence of cognitive capabilities. Postnatal day 24 emerges as a pivotal developmental milestone, marked by the simultaneous stabilization of functional connectivity, refinement of spatial coding, and the emergence of internally-recurring sequences. The developmental shift from encoding elapsed time to encoding distance, likely reflects the progressive integration of extrinsic environmental signals onto an egocentric reference frame. Collectively, these findings indicate that the third postnatal week may constitute a vulnerable developmental period for the emergence of the hippocampal cognitive map.

## STAR Methods

### Mice

All experiments were performed according to the guidelines of the French National Ethics Committee for Sciences and Health report on ‘Ethical Principles for Animal Experimentation’ following European Community Directive 86/609/EEC (Apafis #30.716). We used mouse pups from breedings of Gad-Cre or SST-Cre with wild-type SWISS mice (C.E Janvier, France). All efforts were made to minimize the suffering and the number of animals used.

### Experimental procedures and data acquisition

#### Viruses

*In vivo* calcium imaging experiments were performed using AAV1-hSyn-GCaMP6s.WPRE.SV40 (pAAV.Syn.GCaMP6s.WPRE.SV40 [Addgene viral prep# 100843-AAV1; http://n2t.net/addgene:100843; RRID:Addgene_100843]) and AAV9-FLEX-tdTomato (pAAV-FLEX-tdTomato [Addgene viral prep# 28306-AAV9; http://n2t.net/addgene:28306; RRID:Addgene_28306]).

#### Intracerebroventricular injection

To perform *in vivo* calcium imaging experiments, we first injected viruses into the left lateral ventricle, as previously ^19^. Briefly, newborn GAD1Cre/+ or SST-Cre/+ mice (P0) were anesthetized for 3-4 minutes using hypothermia and injected with 2 µL of a viral mixture consisting of 1.3 µL of AAV2.1-hSynGCaMP6s.WPRE.SV40 and 0.7 µL of AAV9-FLEX-CAG-tdTomato. Blue fast dye (1:20) was added to the mixture to confirm the correct injection by visualizing the ventricle. The coordinates for the left ventricle were visually determined at two-fifths of an imaginary line between the lambda and the left eye, with a depth of 0.4 mm.

#### Surgery for CA1 in vivo 2-photon imaging

The surgery, consisting of implanting a cranial window with a 3 mm diameter above the corpus callosum, was performed as previously described ^50^. Briefly, after being induced using 3% isoflurane in a mix of 90% O2-10% air, P15-16 mice were maintained under 1-3% isoflurane. Analgesia and local anesthesia were ensured by subcutaneous injection of, respectively, buprenorphine (0.1 mg/kg) and lurocaïne. Body temperature was monitored and maintained at 36°C. After securing a headplate (200-200 500 2134-1-5, Luigs & Neumann), the skull and the meninges were removed and the cortex was carefully aspirated until the external capsule became visible. Once the cortectomy was complete, a 3 mm-diameter circular glass coverslip (#1 thickness, Warner Instruments), mounted on a 3 mm-diameter, 1.2 mm-high cannula (Microgroup INC), was sealed in place using KwikSil adhesive (WPI), and the edge of the cannula was secured to the skull with cyanoacrylate. Animals were placed on a heated pad for at least 1 hour after surgery before being returned to their mother and litter. Mice received Carprofen (10 mg/kg) on the day of surgery and for the next three days, and their well-being was carefully monitored.

#### Imaging and behavioral recordings

*In vivo* two-photon calcium imaging experiments were adapted from previous methods ^11^. The animals were head-fixed on a non-motorized treadmill in complete darkness, with no rewards provided, allowing for self-paced locomotion. After 3 or 4 days of habituation, mice were alert but calm and alternated between periods of locomotion and rest during imaging. Mice were divided into two groups: 16 mice were recorded on a 180 or 192-cm belt with no external markers, while 9 mice were recorded on a 192 cm belt enriched in tactile cues. The cued belt was divided into four zones of 48 cm, displaying the following items: (1) paintbrush bristles to stimulate the whiskers; (2) 3.2 mm and 10 mm diameter heat-shrink tubing; (3) 22 mm wide disks of Velcro; (4) cue-poor zone. Treadmill movement was monitored using two alternative systems. In experiments using the uncued 180 cm belt, two pairs of LEDs and photodetectors read periodic patterns on a disk attached to one of the treadmill wheels ^11^. For all experiments with a 192-cm belt, a commercially available treadmill enabled movement and absolute position tracking of the mouse (700-100 100 0012, Luigs & Neumann).

Two-photon calcium imaging was performed using a single-beam multiphoton pulsed laser scanning system coupled to either (1) a TriM Scope II microscope (LaVision Biotech) or (2) an Ultima 2P microscope (Bruker). In both setups, excitation was provided by a Ti:sapphire laser (Chameleon Ultra II, Coherent) tuned to 920 nm for GcaMP and to 1,030 nm for tdTomato excitation. GCaMP and tdTomato fluorescence was isolated using bandpass filter: (1) 510/25 and 580/20 nm, respectively, for the TriM Scope II and (2) 525/25 and 600/20 nm, respectively, for the Ultima 2P. Images were acquired using two GaSP PMTs (H7422-40, Hamamatsu) with a 16x immersion objective (Nikon, NA 0.8). On the TriM Scope II microscope (1), deep and superficial dorsal CA1 pyramidal cell layer fields were imaged in separate sessions. Acquisition was controlled using Imspector software (LaVision Biotech), with a 400 × 400 μm field of view sampled at 9 Hz and a dwell time of 1.85 μs per pixel (2 μm/pixel). On the Ultima 2P microscope (2), deep and superficial CA1 regions were imaged simultaneously using Prairie View software. Using an eletrotunable lens (ETL) and a resonant scanner, the two fields of view (400 × 400 μm; 256 × 256 pixels each) were acquired at 15-16 Hz per plane. In both configurations, the deep and superficial fields of view were spaced 30 μm apart, and each imaging session lasted 20 minutes. To ensure cell tracking across sessions, the alignment of the head mount relative to the microscope was maintained, and the imaging plane was manually adjusted in x, y, and z to match reference tdTomato images acquired on the first imaging day.

Mouvement, position, and image triggers were synchronously acquired and digitized using a 1440A Digidata (Axon instrument, 50 kHz sampling) and the Axoscope 10 software (Axon instrument).

A typical imaging session occurred as follows: (1) the mouse was handled, then fixed on the treadmill; (2) CA1 activity recording for 20 minutes; (3) the mouse returned to its homecage. The next imaging session occurred 24 hours after and mice were imaged for up to 8 consecutive days.

#### Data pre-processing

Image series were motion corrected using either the NoRMCorre algorithm available in the CaImAn toolbox ^51^ or the motion correction included in the Suite2P imaging processing pipeline52.

Cell segmentation was achieved using Suite2p ^52^, based on activity (tau: 1.5 ms, equivalent to the GCaMP6s time constant). To ensure correct segmentation of somatic calcium activity, the automatic detection was manually refined by adding and removing regions of interest (ROIs) with visual inspection of mean projections and fluorescence traces.

For analysis, all of the preprocessed datasets were combined into a single NWB:N ^53^ file per imaging session containing the imaging data, cell contours, animal speed and position, and metadata.

### Data analysis

#### Behavioral analysis

Behavioral data from sessions in which mice completed at least 5 laps and exhibited sufficient running activity (≥8 running periods with speeds >2 cm/s) were analyzed in MATLAB (MathWorks).

Positions were normalized to belt length (192 or 180 cm) and stop positions were identified as the location where a running epoch ended, provided that the preceding run covered at least 10 cm. Cumulative distributions of stop positions were computed per treadmill condition (cued vs. uncued) and statistical comparisons were performed using the Kolmogorov-Smirnov test against a uniform distribution.

#### Calcium event detection and spatial information

Calcium events were detected from DF/F0 fluorescence traces using a smoothed peak-detection algorithm. For each neuron, traces were smoothed with a Gaussian filter (sigma = 5 frames), and peaks were identified as local maxima exceeding the neuron’s median plus one standard deviation, with a minimum prominence proportional to the standard deviation and a minimum inter-event interval of 1 s. Event rates were computed as the number of detected events divided by the recording duration (events/min) for each neuron. Spatial information was calculated as the Shannon information between binned positional occupancy and the neuron’s mean activity per spatial bin, using 32 spatial bins and averaging per bin.

#### Place cells detection

To identify place cells, fluorescence traces were first normalized by z-scoring. Only running periods from mice that completed at least 5 entire laps with sufficient running activity were considered. Treadmill position was normalized to belt length and divided into 100 spatial bins and mean z-scored fluorescence was computed for each bin across laps. To establish a statistical threshold for each cell, we performed 1000 circular shuffles in which the fluorescence signal for each individual lap was randomly shifted, then averaged across shuffled laps and the maximum peak was extracted. The 99th percentile of these 1000 peak values served as our cell-specific statistical threshold. Finally, a cell was classified as a place cell if its actual activity profile contained peaks that exceeded this statistical threshold, had a prominence greater than the threshold value and showed a width between 5 and 30 bins. This approach ensures that identified place fields are both statistically significant and have biologically realistic spatial dimensions.

#### Place field fitting and classification

Since mice were running on a circular track, we fitted place fields using a von Mises distribution, which is a circular analogue of the normal/Gaussian distribution (used conventionally when fitting place fields on linear tracks or in 2D environments ^54,55^, see Methods). Fitting place fields allowed us to estimate their center location and width for each cell on each day, as well as to determine whether the cell was better fit with a single (Figure 2A1, bottom panel) or a double place field model (Figure 2A2, bottom panel; Figure S2A). Neuronal spatial tuning was quantified by fitting place fields to z-scored, binned firing rate traces using a mixture of two von Mises distributions. The model was defined as:

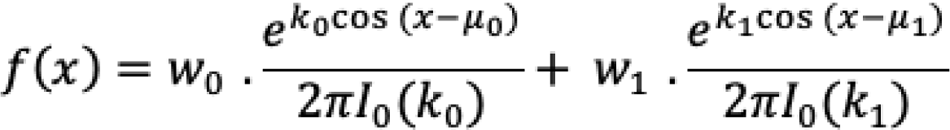

where *x* represents position along the circular environment, μ_0_ and μ_1_ are the mean directions (centers of the distributions), κ_0_ and κ_1_ are concentration parameters (controlling the dispersion of the distribution, where for moderate to large values of κ, the variance can be approximated as σ2≈1/κ), w_0_ and w_1_ are the relative weights of the two components, and I_0_ is the modified Bessel function of order 0.

Fitting was performed using non-linear least squares optimization to identify the parameters that best described each cell’s place field. The fitted mixture model was then decomposed into its two von Mises components, f_0_(*x*) and f_1_(*x*), which were summed to obtain the total fitted function f_total_(*x*)=f_0_(*x*) + f_1_(*x*).

To evaluate the quality of the fit and determine whether a cell exhibited a single or a double place field, we quantified how well each von Mises component accounted for the total fitted function at its center. Specifically, the ratios f_0_(μ_0_)/f_total_(μ_0_) and f_1_(μ_1_)/f_total_(μ_1_) were computed. High ratios (both > 0.9) indicated that each component contributed substantially and independently to the overall fit, suggesting the presence of two distinct components. Among these, the fitted field was classified as double-peaked only if the component with the smaller amplitude reached at least 20 % of the amplitude of the larger component; otherwise, it was classified as single-peaked. This procedure ensured that double-peaked fields reflected genuinely distinct and well-fitted subcomponents, rather than artifacts or poorly separated fits.

#### AUC calculation for place fields

To assess the relative contribution of each neuron across different zones of the circular environment (treadmill), the area under the curve (AUC) of the place field fits was computed within predefined spatial zones (see fig 1C, cued treadmill). Each place field fit was divided into four equally sized zones, and the AUC within each zone was calculated using numerical integration (trapezoidal rule).

#### Place field width estimation (FWHM)

To quantify the spatial extent of each neuron’s place field, the full width at half maximum (FWHM) of the place field fit was calculated. For each neuron, the half-maximum was defined as the midpoint between the minimum and maximum values of the fitted field. The positions at which the fit crossed this half-maximum were identified using linear interpolation between adjacent bins to improve spatial precision. The distance between these positions was taken as the FWHM, providing a measure of the width of the place field.

#### Sequences

We detected neuronal sequences using a method adapted from ^11^. First, we applied offset Principal Component Analysis (PCA ^56^) to smoothed GCaMP fluorescence traces from detected cells and selected the first 5 components. For each component, we cross-correlated its first derivative with individual neuronal traces. Neurons showing correlation values above a threshold determined by Otsu’s method ^57^ were classified as sequence-participating cells. These neurons were then sorted by their time lag of maximal cross-correlation.

Since the first principal components often capture synchronous activation patterns, we excluded components where more than 30% of participating neurons fell within a single 500 ms time bin, as these likely reflect synchronous rather than sequential activity. To validate our detection method and select the optimal component, we manually annotated sequence onset and offset times.

For each manually labeled sequence, we determined the “local order” by cross-correlating individual neuronal activity with the principal component’s first derivative from onset to offset. We then calculated Kendall’s tau rank correlation between this local order and the previously identified neuronal sequence, generating a distribution of correlation values for each component across sequence repetitions.

We selected the principal component with the highest median Kendall’s tau value and a median p-value below 0.05. An imaging session was considered to contain sequences if at least one component met these statistical criteria and involved more than 20 neurons.

For each session, we assessed whether sequence dynamics depended on run distance or run duration by computing Spearman correlations between median running speed and sequence slopes. First, the number of active neurons among the sequence-participating cells was calculated per individual sequence. For each sequence-participating cell, ΔF/F was calculated using the 8th percentile within a sliding window of 1,000 frames and significant calcium transients were identified ^58^. A cell was considered active in an individual sequence if a significant transient occurred between the onset and offset of that sequence. Temporal slope was defined as the ratio of the number of active cells per sequence to sequence duration, and distance slope as the ratio of sequence distance to the number of active cells. Correlations with p < 0.05 were considered significant. Correlation values were then grouped by age (P20-P30) to assess developmental trends using Spearman correlation.

#### SCE detection

For each cell, a 3rd-order polynomial was fit to its time series and the trace was divided by the polynomial fit (per-cell normalization of slow decay). We then applied a Savitzky-Golay filter (order 3, 7-frame window, ∼450 ms) to reduce high-frequency noise and normalized each trace by its median to make amplitudes comparable across cells. Locomotion speed defined behavioral states, with rest as speed ≤ 2 cm/s and run as speed > 2 cm/s, and events detected during run were excluded from all analyses. Calcium transients were detected on the preprocessed traces with the matlab function “findpeaks” using a per-cell prominence threshold of 3 interquartile range and a minimal inter-event interval of 3 frames; Synchronous activity was quantified with a sliding window of length ∼200 ms, by counting the number of distinct cells active within each window. SCE were defined as local maxima of this summed signal that recruited at least 10 cells, with a minimal inter-SCE interval of 5 frames (∼320 ms). For each detected SCE at frame ttt, cell participation was defined as the elementwise maximum of the raster from t−1 and t+2, yielding a binary participation matrix (cells by SCE).

#### Identification of cell assemblies in SCEs

Cell assemblies were identified by clustering SCEs based on the similarity of their cell-participation profiles, followed by statistical testing of cell participation within each SCE cluster. Pairwise SCE dissimilarities were computed as the squared Euclidean distance between columns of the normalized covariance matrix, which improved clustering performance. Unsupervised clustering was performed with k-means at k=10; we ran 2000 replicates and retained the solution with the highest average silhouette value, using MATLAB’s kmeans with ‘plus’ initialization (k-means+), MaxIter=300, and OnlinePhase=’on’ to consolidate solutions.

For an element i, the silhouette was s=(b−a)/max⁡(a,b), where a is the mean dissimilarity of i to its own cluster and b is the lowest mean dissimilarity to any other cluster; here, dissimilarity was derived from the normalized covariance. A null distribution of average silhouette values was obtained by reshuffling cell participation across SCEs and repeating the same procedure; clusters whose average silhouette exceeded the 95th percentile of the null were deemed significant. Each significant SCE cluster was mapped to a cell assembly comprising cells whose participation in that cluster exceeded the 95th percentile of reshuffled data (fraction of SCEs in the cluster that activated the cell). If a cell was significant for multiple clusters, it was assigned to the one with the highest participation fraction. Assembly overlap was quantified per cell using a silhouette based on the normalized Hamming distance between cell pairs; a cell was considered uniquely assigned if this silhouette exceeded the 95th percentile of the reshuffled distribution. Finally, SCEs were ordered by their projection onto assemblies, and an SCE was said to activate an assembly when the number of recruited cells in that assembly exceeded the 95th percentile of chance.

#### Resting correlations of sequence-participating neurons

We tested whether cells involved in activation sequences at J+1 exhibited, on the previous day (J0), spontaneous activity that was more correlated than expected by chance. Only session pairs in which neuronal assemblies were detected at J0 and internal sequences were identified at J+1 were included in this analysis. Cells of interest were identified as those that, on day J0, corresponded (via Track2p) to cells participating in an internal sequence detected at J+1. For each imaging plane, z-scored traces from these cells were extracted during rest periods recorded on J0. Pearson correlation was computed for all possible pairs of these cells and the mean of these values was used as a global measure of co-activation at rest.

To assess whether this mean correlation differed from chance, a non-parametric permutation test was performed. At each iteration, a random set of cells of identical size was drawn from the total population (excluding the cells involved in the sequence at day J+1) and the mean correlation between all pairs in this set was calculated. This procedure was repeated 1 000 times, generating an empirical distribution of mean correlations under the null hypothesis (H₀: no specific co-activation).

The p-value was calculated as the proportion of permutations for which the mean correlation was greater than or equal to the observed mean:

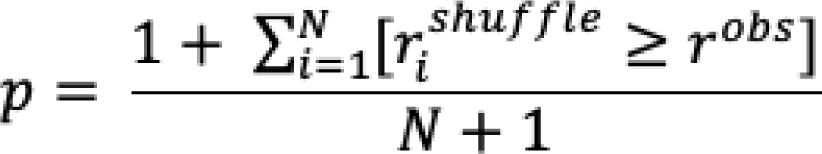

where r^obs^ is the mean correlation among the sequential cells, r_i_^shuffle^ is the mean correlation for the i-th permutation, and N is the total number of permutations. The addition of 1 to both the numerator and denominator is a standard continuity correction to avoid zero p-values.

#### Pairwise Correlation of Place Fields in Pyramidal Cells Across Days

To assess the consistency of place fields over time, we computed the correlation of place fields (*fit*, see *Place field fitting and classification* in Data Analysis) for all possible pairs of imaging days from P20 to P27. Only cells identified as place cells on both days of a given pair were included. For each cell present on both sessions, the Pearson correlation was calculated between its fitted place fields.

#### Place cells anatomical-place field correlations and assemblies topological distribution

For the cued place cell dataset, pairwise anatomical distances between identified place cells were compared to pairwise distances between their place field peaks along the treadmill track. Place field distances were calculated as circular distances on the normalized track length. Correlations between anatomical and functional distances were quantified using Spearman’s correlation. Group-level summaries were obtained by pooling the Spearman correlation values between anatomical and place field distances from all sessions within each age group (P20 −21, P22-23, P24-25, P26-27).

For the internal sequences (uncued conditions) and assemblies (both conditions) datasets, topological analyses were applied to quantify the spatial organization of cells, following previously described approaches ^13,59^. For each cluster (assembly or sequence), the mean silhouette value was computed using pairwise Euclidean distances between cell centroids as a dissimilarity metric. Statistical significance was assessed by comparing the empirical silhouette to a distribution obtained from 1000 random permutations of cell-to-cluster assignments. Spatial clustering was considered significant if the empirical value exceeded the (1 - 0.05/number of clusters) percentile of the shuffled distribution, corresponding to a Bonferroni correction for multiple comparisons.

#### PCA embeddings of neural activity

To visualise the topology, organisation and stability of population activity in the cued conditions we embedded neural activity in PCA space and visualised it as a scatter plot in two dimensions. To visualise the correspondence of the neural state space with the physical space on the treadmill we color-coded each point by the instantaneous location of the mouse at that time point. The color coding was done independently, with the embedding being fully unsupervised, without access to position data. When visualising the cross-day stability of topology and representation we performed the same analysis but fitting PCA on one day and projecting each of the other recordings to that space. Sklearn ^60^ was used to perform the PCA analysis described here.

#### Decoding

All decoding was done using linear regression with ridge regularisation (ridge regression) to avoid overfitting given the large number of neurons, similarly as described in ^29^. Ridge regression optimises the weights (β) that minimise the following loss function:

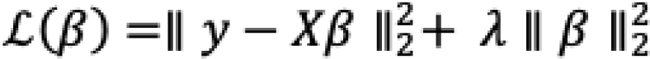

Where in our case y is the position of the mouse on the treadmill, X is neural data and λ is a regularisation parameter. Since mice were running on a treadmill we interpreted the position as an angular variable (0 to 2π) and represented it using polar coordinates when performing the fitting and inference. We also used this representation to compute the R^2^ values, taking into account the circularity of the prediction. For visualisation purposes (Figure 3C, Figure S3B) the prediction of the model was transformed back to the range of the treadmill (0 to 192 cm). To choose the optimal λ and to reliably estimate the model performance on same-day decoding we used nested cross-validation as described previously ^29^. For cross-day decoding we fit a new model with the optimal λ for that given day and evaluated it on all other days. All models were implemented and fitted using PyTorch (Python).

#### Hippocampal Network functional connectivity

To compute the functional connectivity, we started from the z-scored fluorescence traces during running activity (speed > 2 cm/s). Pairwise Pearson correlations were computed between the activity traces of all neurons. Statistical significance was assessed using a block-shuffling permutation test that preserves local temporal correlations. Time-series were divided into contiguous 15 seconds blocks, which were randomly permuted across n = 1000 repetitions to generate a null distribution of correlation values. For each neuron pair, the empirical correlation was compared against this null distribution, and only values exceeding a two-tailed significance threshold (α=0.01) were retained for subsequent functional connectivity analysis. Furthermore, to reduce the potential contamination from nearby ROIs or local signal bleed-through, we conservatively excluded all functional connections between different layers that were separated by less than 7.8 µm (5 pixel) in the field of view, based on the histogram of pairwise distances.

The Functional Connectivity Similarity (FCS) between two days was defined as the Pearson correlation between the weights of the edges across the two days, as previously described ^29^. In the present analysis, only edges present on both days were considered. We fitted a linear mixed-effects model with period (early vs. late) and day difference as fixed effects and mouse as a random intercept (R, lme4). Results were robust to the normalization method, including DF/F0 versus z-scored traces.

## Acknowledgments

We thank all the members of the Cossart lab for helpful discussions and constructive feedback. We thank Dr. Sophie Brustlein and Marie Kurz for their technical support, and Dr. Arthur Godino for critical feedback on the analysis. We thank INMED’s animal facility and PBMC technological platform for excellent technical support. We are grateful to Dr. H. Monyer for providing the Gad-Cre mouse line. We are grateful to Drs. D. Dombeck and J. Climer for sharing analysis codes. This work was supported by the European Research Council under the European Union’s Horizon 2020 research and innovation program grant #951330 (HOPE). E.L., C.F. and M.M. were funded by the ERC. R.F.D. was funded by the “Ministère de l’Enseignement Supérieur, de la Recherche et de l’Innovation” and by the Fondation pour la Recherche Médicale (grant no. FDT202106012824). J.M. is funded by the Fondation Roger de Spoelberch. M.B. was funded by the Fyssen Foundation, the Foundation pour la Recherche Médicale (grant no. SPF20170938593), and by the European Union (Marie Skłodowska-Curie individual fellowship, grant no. 794861—IF-2017). V.D. acknowledges support from the PR[AI]RIE-PSL Fellowship in AI. V.D. and R.M. acknowledge support from the Locomat project (ANR-21-CE16-0037). R.C. is supported by the CNRS. J-C. P. and M.A.P are supported by the INSERM.

## Author contributions

E.L., C.F., M.M. and R.C. designed research. E.L., C.F. and M.C. acquired the data with technical help of M.B. and M.A.P.. E.L., C.F., M.M., R.F.D., V.D., R.M., J.M., J-C.P. performed analysis. R.C., E.L., C.F., M.M., M. B., R.F.D., V.D., wrote the paper.

---

**Figure S1.**
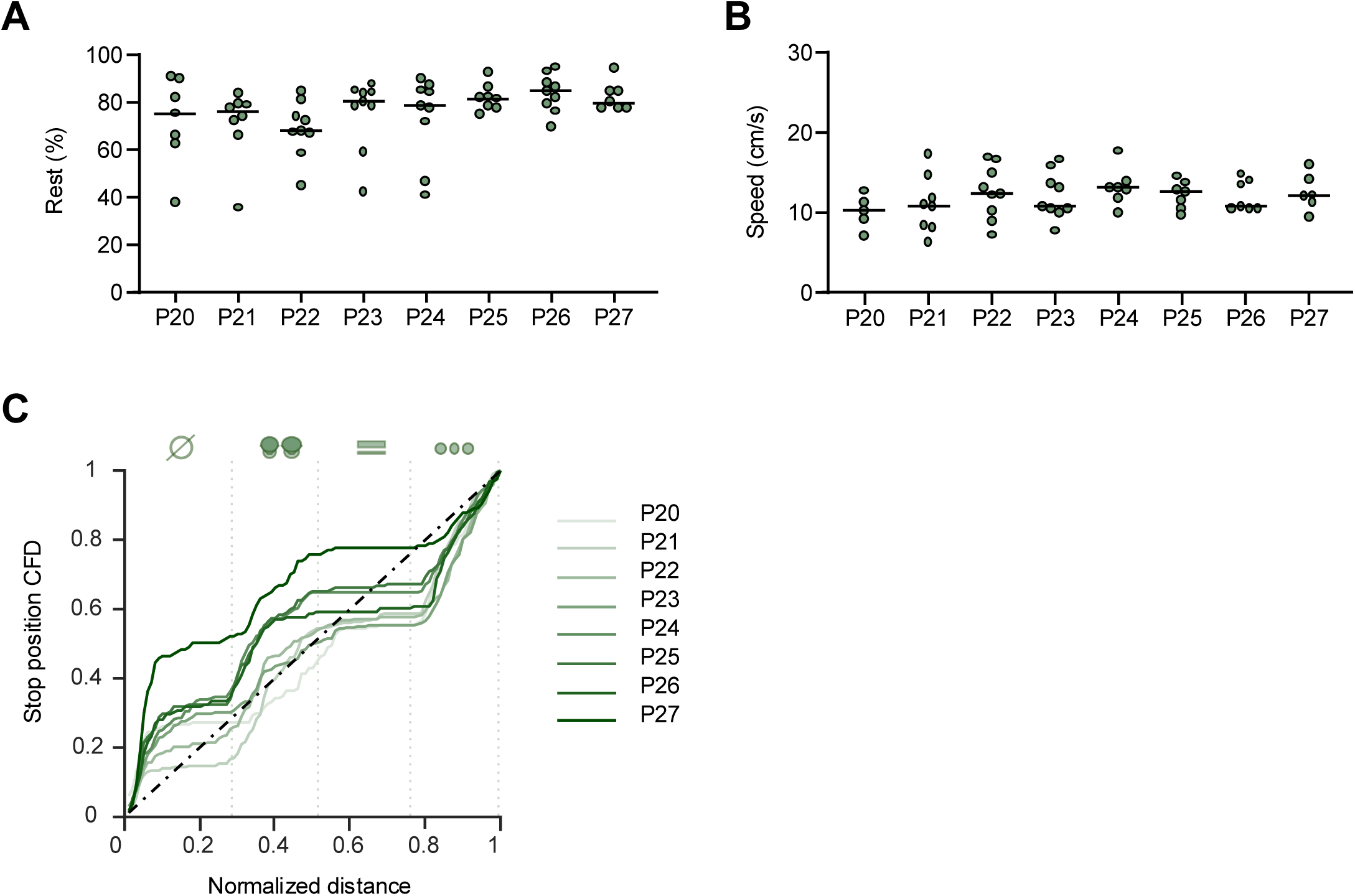
Evolution of the spontaneous behavior of head-fixed mice imaged on cued treadmill during the fourth postnatal week. Related to Figure 1 **(A)** Mean time spent at rest per age on the cued treadmills (P20: median = 75.45 % [iqr = 27.62], N = 7 mice; P21: median = 75.77 % [iqr = 11.72], N = 8 mice; P22: median = 67.60 % [iqr = 14.98], N = 9 mice; P23: median = 80.44 % [iqr = 15.95], N = 9 mice; P24: median = 78.41 % [iqr = 27.16], N = 9 mice; P25: median = 81.75 % [iqr = 7.65], N = 8 mice; P26: median = 84.57 % [iqr = 13.04], N = 9 mice; P27: median = 80.00 % [iqr = 7.05], N = 7 mice). No significant change is observed across ages (LMM F_(1,7)_ = 1.76, p = 0.12). Each dot represents the mean rest time per animal per age. Black lines indicate median values. **(B)** Mean running speed on the cued per age on the cued treadmills (P20: median = 10.26 cm/s [iqr = 3.90], N = 5 mice; P21: median = 10.86 cm/s [iqr = 5.77], N = 8 mice; P22: median = 12.27 cm/s [iqr = 6.27], N = 9 mice; P23: median = 10.74 cm/s [iqr = 4.49], N = 9 mice; P24: median = 13.12 cm/s [iqr = 2.05], N = 7 mice; P25: median = 12.68 cm/s [iqr = 3.16], N = 7 mice; P26: median = 10.77 cm/s [iqr = 3.58], N = 7 mice; P27: median = 12.16 cm/s [iqr = 3.85], N = 6 mice). No significant change is observed across ages (LMM F_(1,7)_ = 0.97, p = 0.47). Each dot represents the mean rest time per animal per age. Black lines indicate median values. **(C)** Cumulative frequency distribution (CFD) of stop positions per age along the cued treadmills. CFDs were compared to a uniform distribution (0-1, black dashed line) of matching sample size. The distribution shows preferred stop positions on the cued condition for each age (Kolmogorov-Smirnov test, P20: D = 0.2249, p < 0.0001; P21: D = 0.2020, p < 0.001; P22: D = 0.2083, p < 0.001; P23: D = 0.2359, p < 0.0001; P24: D = 0.1918, p < 0.01; P25: D = 0.2068, p < 0.001; P26: D = 0.2024, p < 0.001; P27: D = 0.3752, p < 0.0001). Each line represents the cumulative distributions from all animals per age.

**Figure S2.**
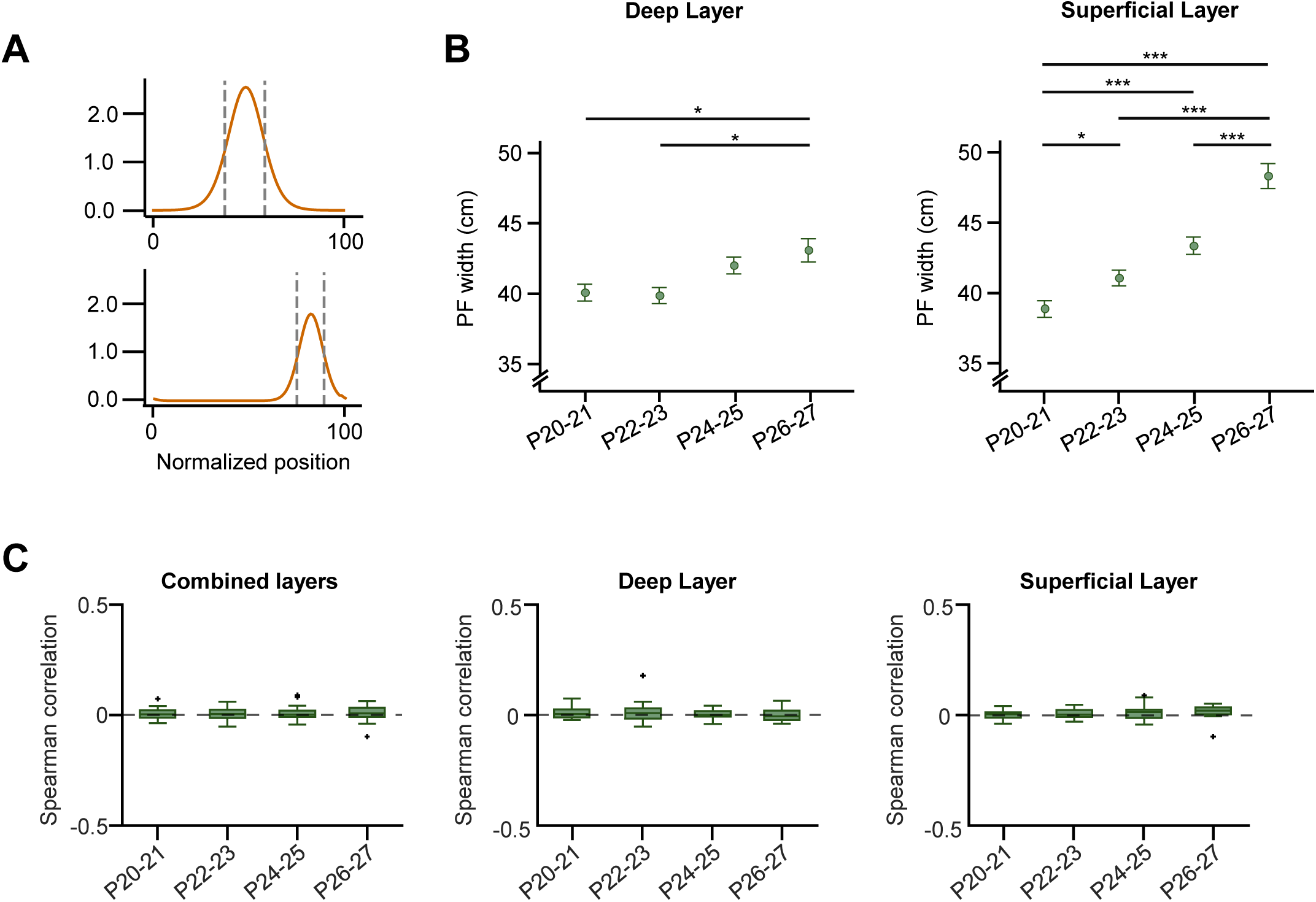
Developmental maturation of CA1 place field properties across hippocampal sublayers. Related to Figure 2. **(A)** Decomposition of the von Mises mixture fits from a neuron with multiple place fields along normalized position, showing the two individual component distributions with their respective full width at half maximum. **(B)** Place field width across age groups for CA1 deep (P20-21: 40 ± 0.6cm n = 1388 PCs N = 7 mice, P22-23: 40± 0.6cm n = 1399 PCs N = 8 mice, P24-25: 42 ± 0.6cm n = 1207 PCs N = 7 mice, P26-27: 43 ± 0.8cm n = 750 PCs N = 7 mice) and superficial sublayers separately (P20-21: 39 ± 0.6cm n = 1535 PCs N = 7 mice, P22-23: 41 ± 0.6cm n = 1707 PCs N = 8 mice, P24-25: 43 ± 0.6cm n = 1682 PCs N = 7 mice, P26-27: 48 ± 0.9cm n = 1101 PCs N = 7 mice; See Table S5 for detailed statistics). Data are presented as mean ± SEM. **(C)** Relationship between CA1 place cells anatomical distance and their place field separation as a fonction of the age in both layers (lelft panel, P20-21: median = 0.005 [iqr = 0.029], P22-23: median = 0.005 [iqr = 0.036], P24-25: median = 0.003 [iqr = 0.028], P26-27: median = 0.009 [iqr = 0.041]), in deep layer (middle panel, P20-21: median = 0.005 [iqr = 0.033], P22-23: median = 0.009 [iqr = 0.045], P24-25: median = 0.001 [iqr = 0.025], P26-27: median = −0.006 [iqr = 0.042]), and in superficial layer (right panel, P20-21: median = 0.005 [iqr = 0.022], P22-23: median = 0.003 [iqr = 0.031], P24-25: median = 0.015 [iqr = 0.038], P26-27: median = 0.021 [iqr = 0.031]). No significant change between the age groups when looking at deep and superficial planes together (LMM, F_(3,92.53)_ = 0.105, p = 0.957) or separately (LMM, age x plane interaction: F_(3,89.17)_ = 0.771, p = 0.514). See Table S7 for detailed statistics. Statistical comparison performed using Linear mixed model (random effect: mouse identity); *p < 0.05, **p < 0.01, ***p < 0.001.

**Figure S3.**
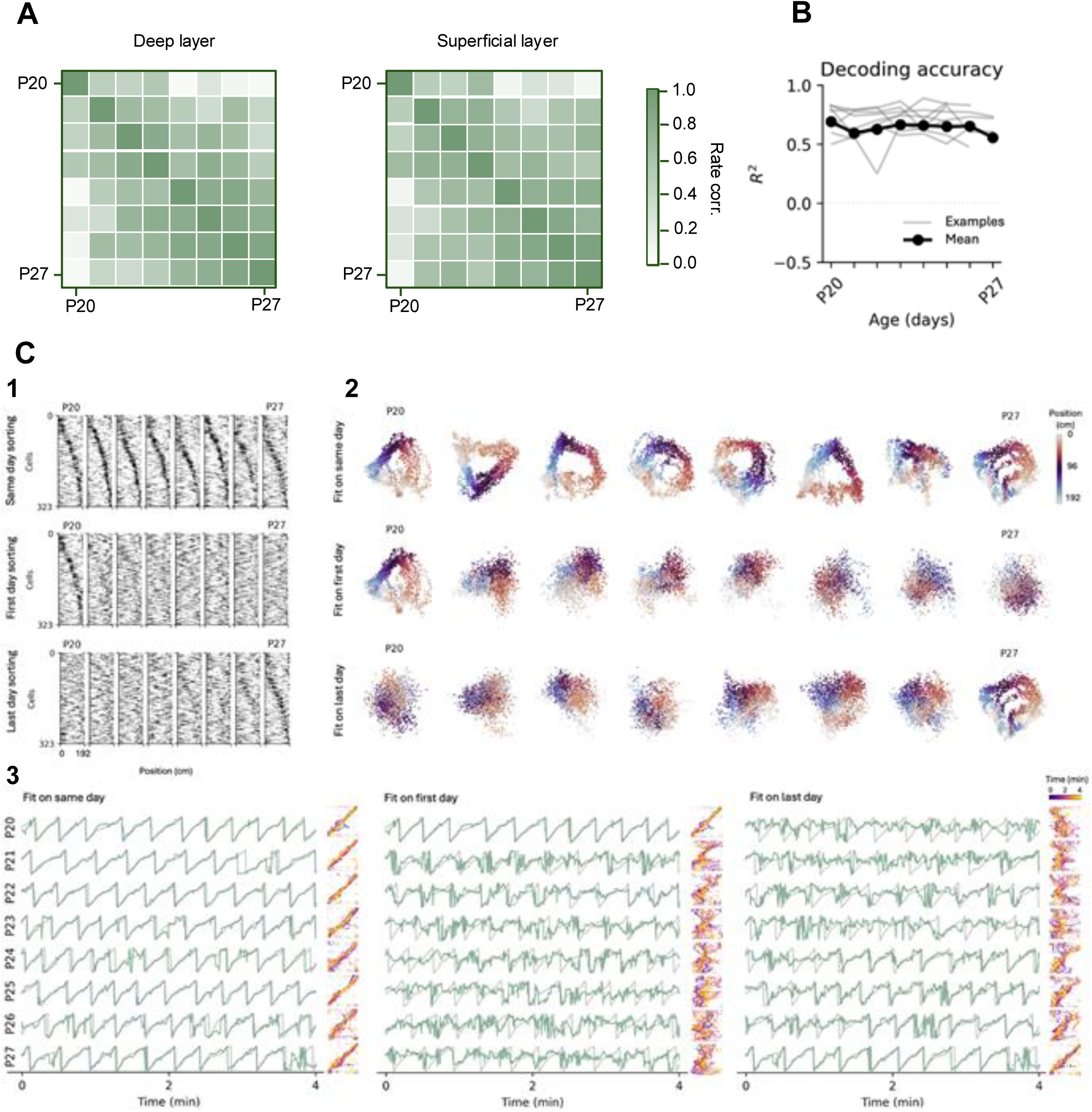
Longitudinal analysis of spatial representation stability in developing CA1. **Related to** Figure 3 **(A)** Mean pearson correlation of placefields for all day pairs averaged across all animals for CA1 deep and superficial sublayers separately (see Table S8 for detailed statistics). **(B)** Coefficient of determination (R^2^) values from same-day decoding of position based on simultaneously recorded neural activity as a function of age. Individual mice are shown in grey and the mean across mice in black. **(C)**(1) Spatially averaged activity of all tracked cells. Top: Sorted by peak position computed on the same day. The sorting was computed on the first half of the data and applied to the second half. Note that the cell correspondence is shuffled here due to individual sorting. Middle: Sorted by peak position on the first recording day (P20). Bottom: Same as middle but sorted on the last day (P27). (2) PCA embeddings of instantaneous neural activity color-coded by the mouse position. Top: PCA was fit individually on each day and data was projected to the PC space of that same day. Middle: PCA was fit on the first day (P20) and activity of all other days was projected to that PC space. Right: Same as middle but fit on the last day (P27). (3) Position decoding from simultaneously recorded neural activity. Left: The fit and decoding are done for the same day using cross-validation. Grey denotes the true position of the mouse and green the prediction of the decoding model. Plots to the right of the time series are a scatter between the true position and the prediction, with values on the identity line denoting good correspondence. Color code corresponds to time. Middle: The fit was done on the first day (P20) and the same model was used to try to decode position across all other days (‘cross-day decoding’). Right: Same as middle but fit on the last day (P27).

**Figure S4.**
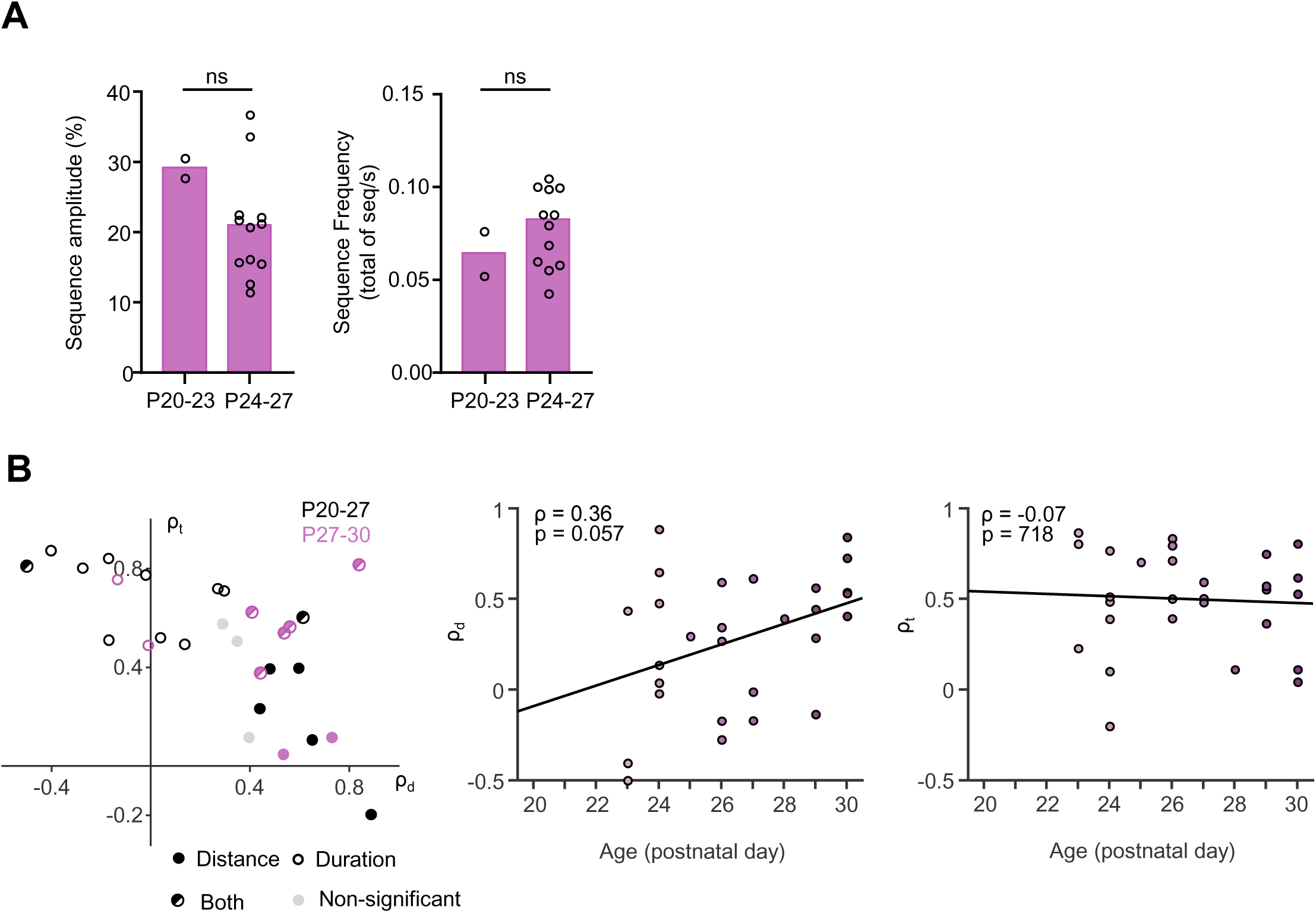
Characterisation of the internal sequences during the fourth postnatal week. Related to Figure 4. Left: Proportion of session displaying internal sequences (pink) in the deep layer et P20-23 (1/13 sessions), P24-27 (9/20 sessions). Empty parts represent the proportion of sessions without internal sequences. *: p = 0.050. Right: Same as Left for superficial layer, with 2/13 sessions at P20-23 and 5/20 sessions at P24-P27. **(B)** Across all imaging sessions (P20-27: n = 17 sessions, 5 mice; P27-30: n = 11 sessions, 3 mice), Spearman correlation coefficients for time (ρₜ) are plotted as a function of Spearman correlation coefficients for distance (ρ_d_). Sperman correlations were calculated considering only significant individual sequences, defined by a significant Kendall’s tau rank correlation with the reference sequence. Sessions recorded between P20-27 and P27-30 are represented by black and pink dots, respectively. Sessions showing significant correlations for distance, time, both, or neither are indicated by filled, open, half-filled, or gray dots, respectively (2) Spearman correlation for time calculated from significant sequences (ρ_t_) as a function of age (P20-P30). Each dot represents an imaging session, and the black line shows a linear fit with Spearman correlation coefficient and p value associated. (3) Same as (2) for Spearman correlation for distance calculated from significant sequences (ρ_d_).

**Table S1.**
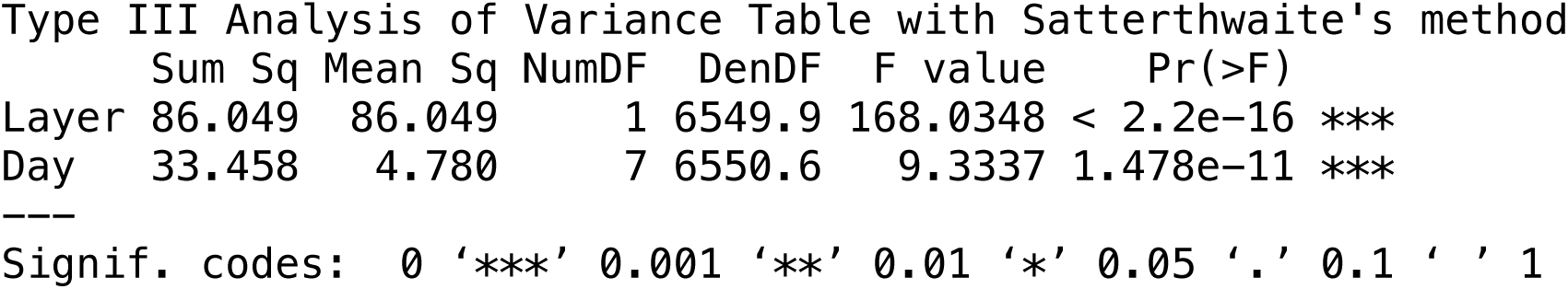

|  | Sum Sq | Mean Sq | NumDF | DenDF | F value | Pr(>F) |
| --- | --- | --- | --- | --- | --- | --- |
| Layer | 86.049 | 86.049 | 1 | 6549.9 | 168.0348 | < 2.2e-16 *** |
| Day | 33.458 | 4.780 | 7 | 6550.6 | 9.3337 | 1.478e-11 *** |

Signif. codes: 0 '\*\*\*' 0.001 '\*\*' 0.01 '\*' 0.05 '.' 0.1 ' ' 1

**Table S2 - Related to Figure 2.**

|  | Sum Sq | Mean Sq | NumDF | DenDF | F value | Pr(>F) |
| --- | --- | --- | --- | --- | --- | --- |
| age | 546.97 | 182.323 | 3 | 89.803 | 3.7427 | 0.01386 * |
| plane | 7.16 | 7.163 | 1 | 89.063 | 0.1470 | 0.70230 |
| age:plane | 82.46 | 27.488 | 3 | 89.063 | 0.5643 | 0.63999 |

--- Signif. codes: 0 '\*\*\*' 0.001 '\*\*' 0.01 '\*' 0.05 '.' 0.1 ' ' 1

**Table S3.**

|  | Sum Sq | Mean Sq | NumDF | DenDF | F value | Pr(>F) |
| --- | --- | --- | --- | --- | --- | --- |
| Layer | 0.74001 | 0.74001 | 1 | 6551.2 | 30.5304 | 3.413e-08 *** |
| Day | 0.67855 | 0.09694 | 7 | 6551.9 | 3.9992 | 0.000225 *** |

Signif. codes: 0 '\*\*\*' 0.001 '\*\*' 0.01 '\*' 0.05 '.' 0.1 ' ' 1

**Table S4 - related to Figure 2.**

|  | Sum Sq | Mean Sq | NumDF | DenDF | F value | Pr(>F) |
| --- | --- | --- | --- | --- | --- | --- |
| age | 235.856 | 78.619 | 3 | 90.698 | 2.3519 | 0.07746 . |
| plane | 166.430 | 166.430 | 1 | 89.356 | 4.9789 | 0.02816 * |
| age:plane | 6.783 | 2.261 | 3 | 89.356 | 0.0676 | 0.97697 |

Signif. codes: 0 '\*\*\*' 0.001 '\*\*' 0.01 '\*' 0.05 '.' 0.1 ' ' 1

**Table S5 - related to Figure 2 and S2.**
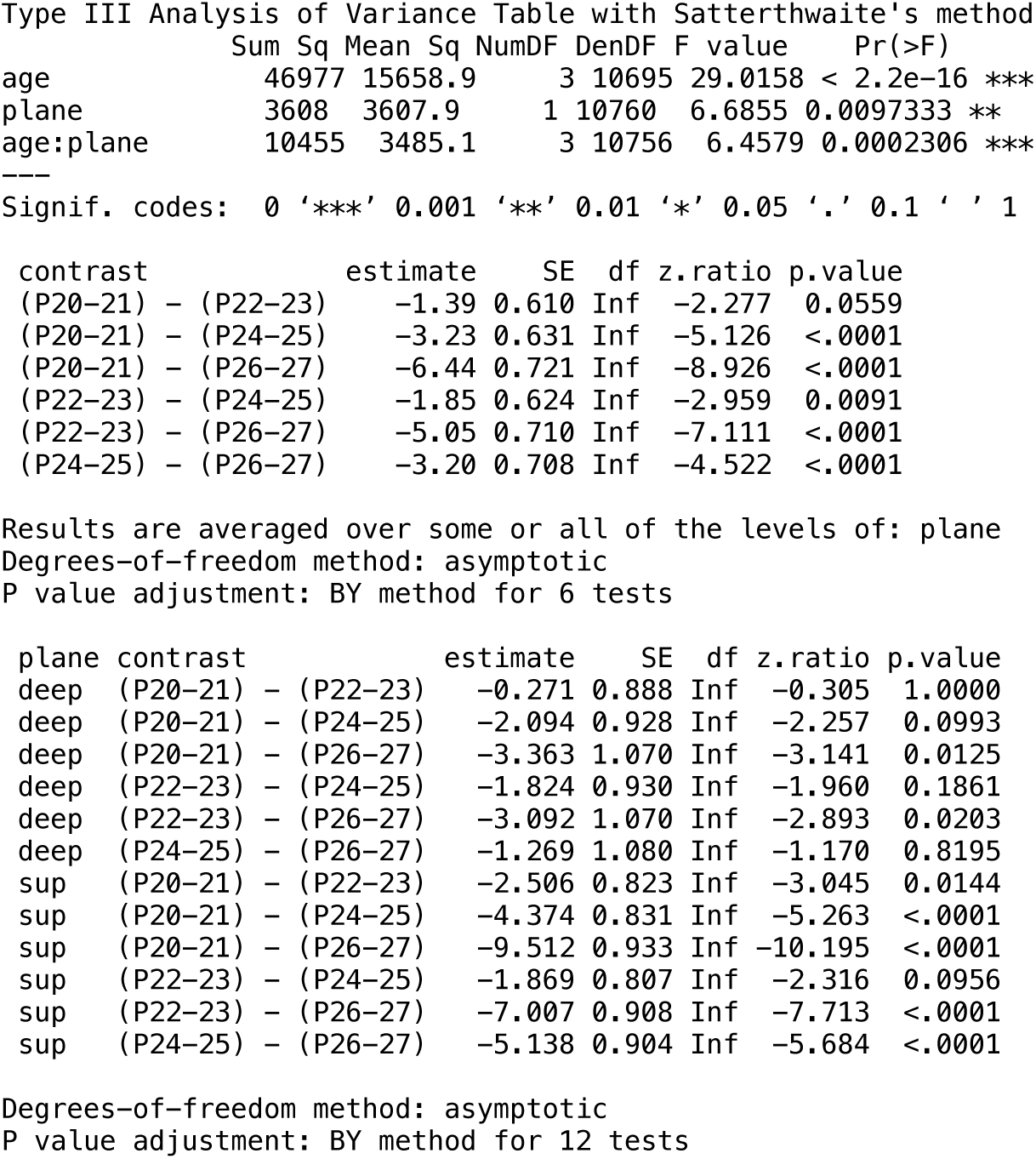

|  | Sum Sq | Mean Sq | NumDF | DenDF | F value | Pr(>F) |
| --- | --- | --- | --- | --- | --- | --- |
| age | 46977 | 15658.9 | 3 | 10695 | 29.0158 | < 2.2e-16 *** |
| plane | 3608 | 3607.9 | 1 | 10760 | 6.6855 | 0.0097333 ** |
| age:plane | 10455 | 3485.1 | 3 | 10756 | 6.4579 | 0.0002306 *** |

| contrast | estimate | SE | df | z.ratio | p.value |
| --- | --- | --- | --- | --- | --- |
| (P20-21) - (P22-23) | -1.39 | 0.610 | Inf | -2.277 | 0.0559 |
| (P20-21) - (P24-25) | -3.23 | 0.631 | Inf | -5.126 | <.0001 |
| (P20-21) - (P26-27) | -6.44 | 0.721 | Inf | -8.926 | <.0001 |
| (P22-23) - (P24-25) | -1.85 | 0.624 | Inf | -2.959 | 0.0091 |
| (P22-23) - (P26-27) | -5.05 | 0.710 | Inf | -7.111 | <.0001 |
| (P24-25) - (P26-27) | -3.20 | 0.708 | Inf | -4.522 | <.0001 |

| plane | contrast | estimate | SE | df | z.ratio | p.value |
| --- | --- | --- | --- | --- | --- | --- |
| deep | (P20-21) - (P22-23) | -0.271 | 0.888 | Inf | -0.305 | 1.0000 |
| deep | (P20-21) - (P24-25) | -2.094 | 0.928 | Inf | -2.257 | 0.0993 |
| deep | (P20-21) - (P26-27) | -3.363 | 1.070 | Inf | -3.141 | 0.0125 |
| deep | (P22-23) - (P24-25) | -1.824 | 0.930 | Inf | -1.960 | 0.1861 |
| deep | (P22-23) - (P26-27) | -3.092 | 1.070 | Inf | -2.893 | 0.0203 |
| deep | (P24-25) - (P26-27) | -1.269 | 1.080 | Inf | -1.170 | 0.8195 |
| sup | (P20-21) - (P22-23) | -2.506 | 0.823 | Inf | -3.045 | 0.0144 |
| sup | (P20-21) - (P24-25) | -4.374 | 0.831 | Inf | -5.263 | <.0001 |
| sup | (P20-21) - (P26-27) | -9.512 | 0.933 | Inf | -10.195 | <.0001 |
| sup | (P22-23) - (P24-25) | -1.869 | 0.807 | Inf | -2.316 | 0.0956 |
| sup | (P22-23) - (P26-27) | -7.007 | 0.908 | Inf | -7.713 | <.0001 |
| sup | (P24-25) - (P26-27) | -5.138 | 0.904 | Inf | -5.684 | <.0001 |

Degrees-of-freedom method: asymptotic
P value adjustment: BY method for 12 tests

**Table S6 - related to Figure 2.**

| Type III Analysis of Variance Table with Satterthwaite's method |  |  |  |  |  |  |
| --- | --- | --- | --- | --- | --- | --- |
|  | Sum Sq | Mean Sq | NumDF | DenDF | F value | Pr(>F) |
| age | 71065 | 23688 | 3 | 42923 | 81.1396 | < 2.2e-16 *** |
| plane | 7512 | 7512 | 1 | 43044 | 25.7314 | 3.940e-07 *** |
| zone | 902972 | 300991 | 3 | 43037 | 1030.9883 | < 2.2e-16 *** |
| age:plane | 838 | 279 | 3 | 43040 | 0.9571 | 0.4119 |
| age:zone | 11979 | 1331 | 9 | 43037 | 4.5591 | 4.969e-06 *** |
| plane:zone | 6171 | 2057 | 3 | 43037 | 7.0457 | 9.883e-05 *** |
| age:plane:zone | 2012 | 224 | 9 | 43037 | 0.7659 | 0.6483 |

| contrast | estimate | SE | df | z.ratio | p.value |
| --- | --- | --- | --- | --- | --- |
| Z1 - Z2 | -9.85 | 0.24 | Inf | -40.972 | <.0001 |
| Z1 - Z3 | -11.78 | 0.24 | Inf | -48.992 | <.0001 |
| Z1 - Z4 | -3.64 | 0.24 | Inf | -15.136 | <.0001 |
| Z2 - Z3 | -1.93 | 0.24 | Inf | -8.020 | <.0001 |
| Z2 - Z4 | 6.21 | 0.24 | Inf | 25.835 | <.0001 |
| Z3 - Z4 | 8.14 | 0.24 | Inf | 33.855 | <.0001 |

| contrast | estimate | SE | df | z.ratio | p.value |
| --- | --- | --- | --- | --- | --- |
| (20-21) - (22-23) | -1.753 | 0.224 | Inf | -7.817 | <.0001 |
| (20-21) - (24-25) | -3.211 | 0.232 | Inf | -13.824 | <.0001 |
| (20-21) - (26-27) | -3.377 | 0.266 | Inf | -12.717 | <.0001 |
| (22-23) - (24-25) | -1.457 | 0.230 | Inf | -6.345 | <.0001 |
| (22-23) - (26-27) | -1.623 | 0.261 | Inf | -6.208 | <.0001 |
| (24-25) - (26-27) | -0.166 | 0.261 | Inf | -0.636 | 1.0000 |

| zone | contrast | estimate | SE | df | z.ratio | p.value |
| --- | --- | --- | --- | --- | --- | --- |
| Z1 | (20-21) - (22-23) | -1.002 | 0.443 | Inf | -2.259 | 0.1140 |
| Z1 | (20-21) - (24-25) | -2.769 | 0.455 | Inf | -6.086 | <.0001 |
| Z1 | (20-21) - (26-27) | -4.136 | 0.518 | Inf | -7.986 | <.0001 |
| Z1 | (22-23) - (24-25) | -1.767 | 0.449 | Inf | -3.933 | 0.0005 |
| Z1 | (22-23) - (26-27) | -3.135 | 0.512 | Inf | -6.121 | <.0001 |
| Z1 | (24-25) - (26-27) | -1.368 | 0.518 | Inf | -2.639 | 0.0470 |
| Z2 | (20-21) - (22-23) | -1.020 | 0.443 | Inf | -2.300 | 0.1080 |
| Z2 | (20-21) - (24-25) | -1.847 | 0.455 | Inf | -4.060 | 0.0004 |
| Z2 | (20-21) - (26-27) | -3.085 | 0.518 | Inf | -5.955 | <.0001 |
| Z2 | (22-23) - (24-25) | -0.827 | 0.449 | Inf | -1.841 | 0.2832 |
| Z2 | (22-23) - (26-27) | -2.065 | 0.512 | Inf | -4.032 | 0.0004 |
| Z2 | (24-25) - (26-27) | -1.238 | 0.518 | Inf | -2.389 | 0.0901 |
| Z3 | (20-21) - (22-23) | -3.235 | 0.443 | Inf | -7.296 | <.0001 |
| Z3 | (20-21) - (24-25) | -4.529 | 0.455 | Inf | -9.955 | <.0001 |
| Z3 | (20-21) - (26-27) | -3.680 | 0.518 | Inf | -7.105 | <.0001 |
| Z3 | (22-23) - (24-25) | -1.294 | 0.449 | Inf | -2.880 | 0.0240 |
| Z3 | (22-23) - (26-27) | -0.445 | 0.512 | Inf | -0.869 | 1.0000 |
| Z3 | (24-25) - (26-27) | 0.849 | 0.518 | Inf | 1.638 | 0.3997 |
| Z4 | (20-21) - (22-23) | -1.757 | 0.443 | Inf | -3.963 | 0.0005 |
| Z4 | (20-21) - (24-25) | -3.699 | 0.455 | Inf | -8.131 | <.0001 |
| Z4 | (20-21) - (26-27) | -2.605 | 0.518 | Inf | -5.029 | <.0001 |
| Z4 | (22-23) - (24-25) | -1.942 | 0.449 | Inf | -4.321 | 0.0001 |
| Z4 | (22-23) - (26-27) | -0.848 | 0.512 | Inf | -1.655 | 0.3997 |
| Z4 | (24-25) - (26-27) | 1.094 | 0.518 | Inf | 2.111 | 0.1576 |

Results are averaged over some or all of the levels of: plane Degrees-of-freedom method: asymptotic P value adjustment: BY method for 24 tests

**Table S7 - related to Figure S2.**
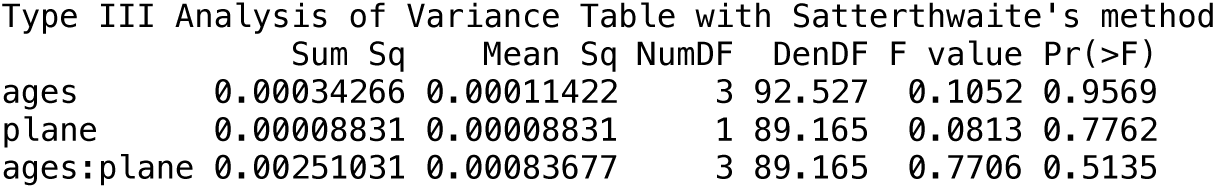

|  | Sum Sq | Mean Sq | NumDF | DenDF | F value | Pr(>F) |
| --- | --- | --- | --- | --- | --- | --- |
| ages | 0.00034266 | 0.00011422 | 3 | 92.527 | 0.1052 | 0.9569 |
| plane | 0.00008831 | 0.00008831 | 1 | 89.165 | 0.0813 | 0.7762 |
| ages:plane | 0.00251031 | 0.00083677 | 3 | 89.165 | 0.7706 | 0.5135 |

**Table S8 - related to Figure 3 and S3.**
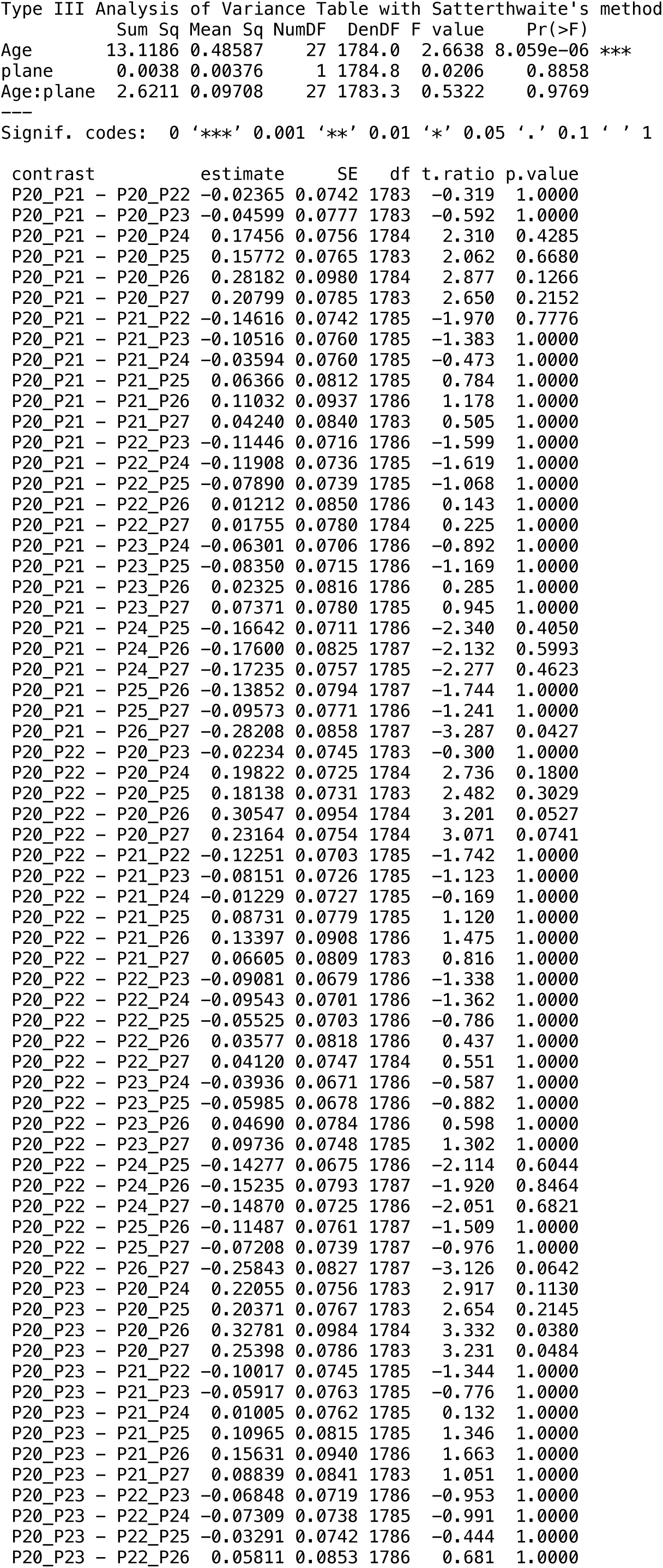
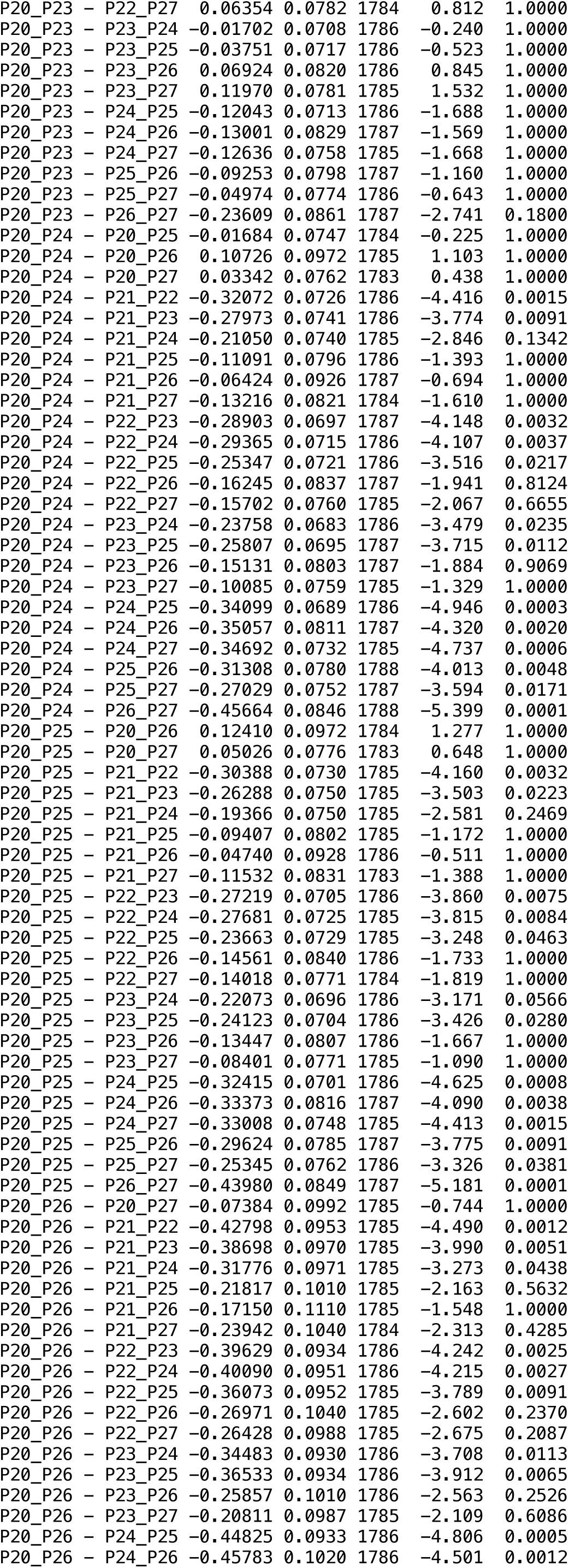
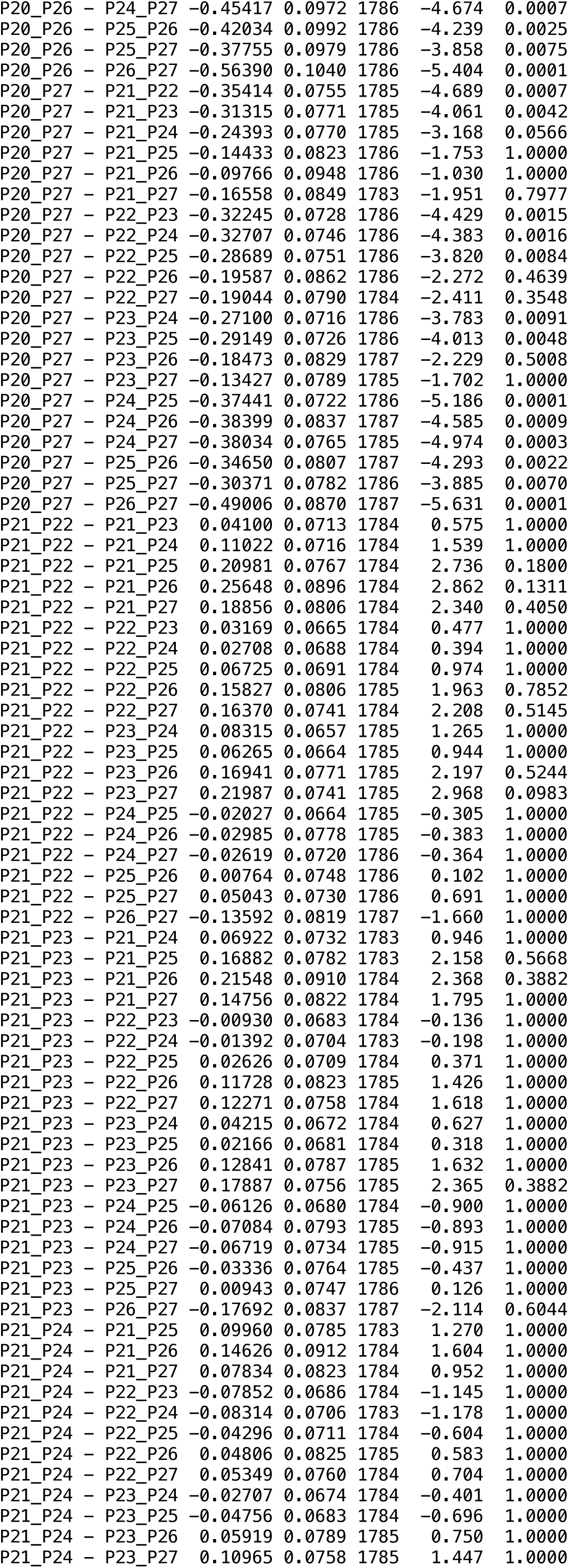
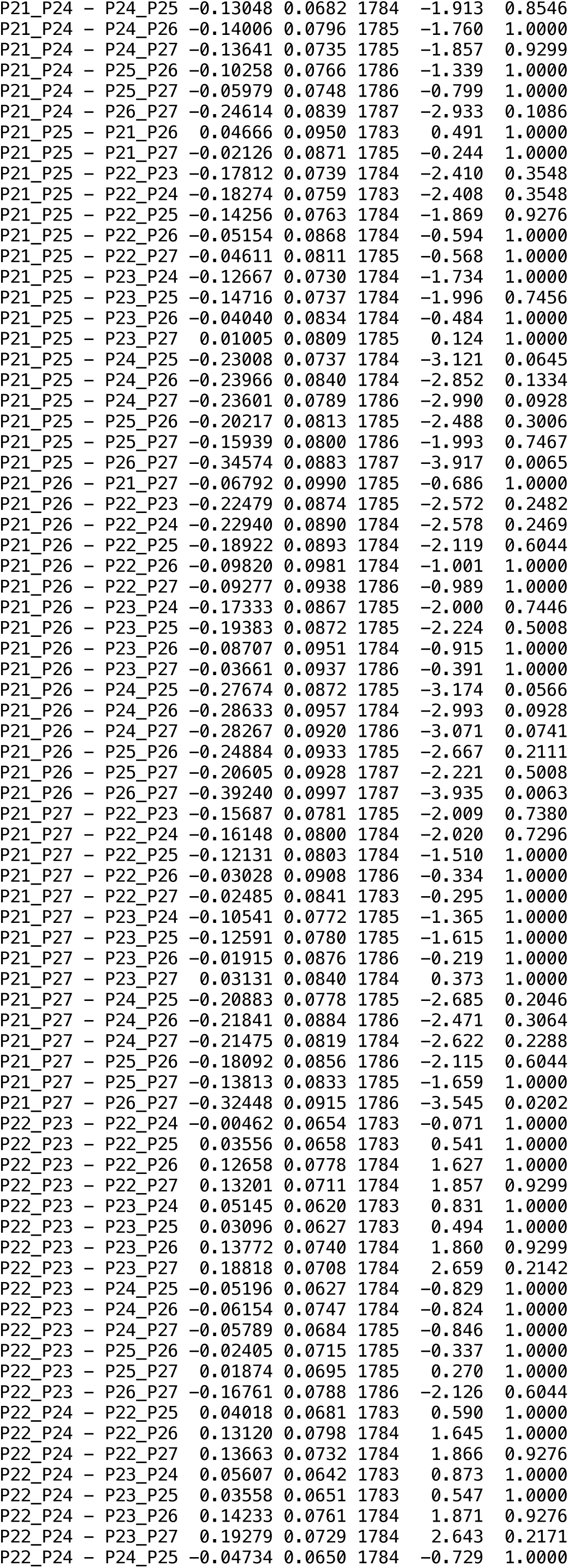
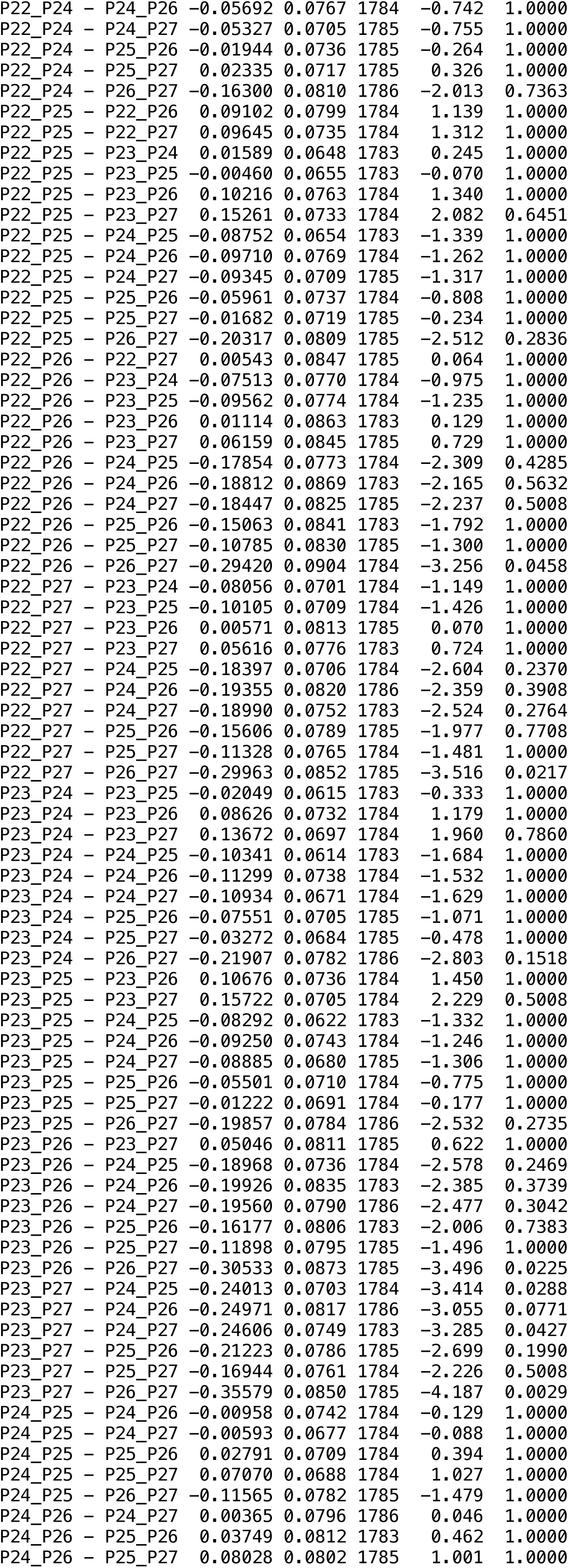
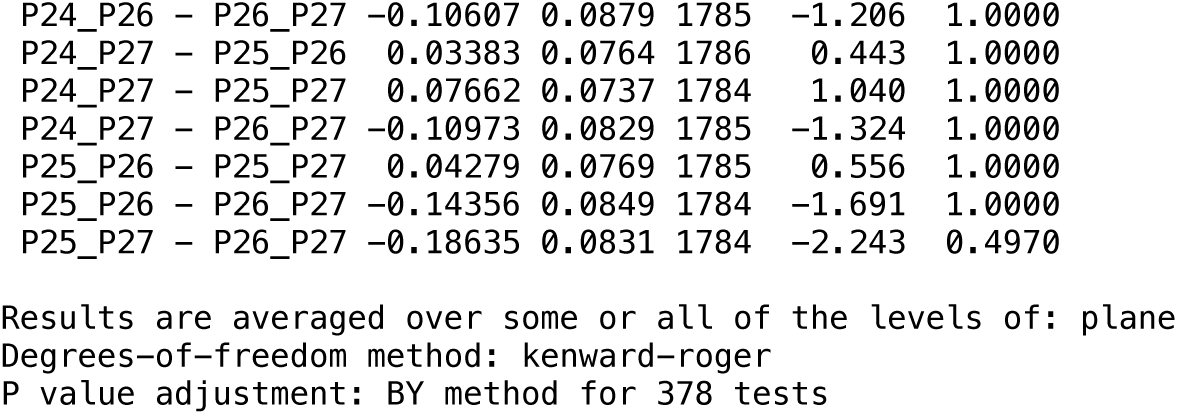

**Table S9 - related to Figure 3.**

|  | Sum Sq | Mean Sq | NumDF | DenDF | F value | Pr(>F) |
| --- | --- | --- | --- | --- | --- | --- |
| period | 0.87350 | 0.87350 | 1 | 1150.5 | 5.3357 | 0.02107 * |
| plane | 0.07250 | 0.07250 | 1 | 1146.9 | 0.4429 | 0.50588 |
| period:plane | 0.02148 | 0.02148 | 1 | 1146.7 | 0.1312 | 0.71722 |

--- Signif. codes: 0 '\*\*\*' 0.001 '\*\*' 0.01 '\*' 0.05 '.' 0.1 ' ' 1

**Table S10 - related to Figure 3.**
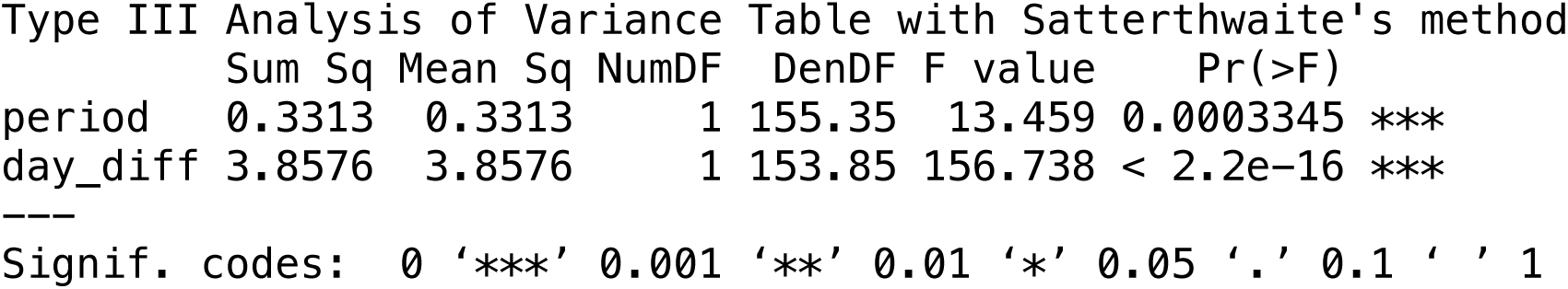

**Table S11 - related to Figure 3.**

|  | Sum Sq | Mean Sq | NumDF | DenDF | F value | Pr(>F) |  |
| --- | --- | --- | --- | --- | --- | --- | --- |
| period | 0.167738 | 0.167738 | 1 | 57.686 | 18.2631 | 7.295e-05 | *** |
| day_diff | 0.014459 | 0.014459 | 1 | 57.995 | 1.5743 | 0.2146 |  |

Signif. codes: 0 '\*\*\*' 0.001 '\*\*' 0.01 '\*' 0.05 '.' 0.1 ' ' 1

**Table S12 - related to Figure 4.**

|  | Chisq | Df | Pr(>Chisq) |  |
| --- | --- | --- | --- | --- |
| (Intercept) | 9.2681 | 1 | 0.002332 | ** |
| period | 6.8678 | 2 | 0.032260 | * |
| plane | 0.3476 | 1 | 0.555489 |  |
| period:plane | 1.2532 | 2 | 0.534405 |  |

| contrast | estimate | SE | df | z.ratio | p.value |
| --- | --- | --- | --- | --- | --- |
| (20-23) - (24-27) | -1.684 | 0.738 | Inf | -2.281 | 0.0338 |
| (20-23) - (27-30) | -2.601 | 0.806 | Inf | -3.227 | 0.0038 |
| (24-27) - (27-30) | -0.916 | 0.637 | Inf | -1.440 | 0.1500 |

Results are averaged over some or all of the levels of: plane
Results are given on the log odds ratio (not the response) scale.
P value adjustment: BH method for 3 tests

**Table S13.**
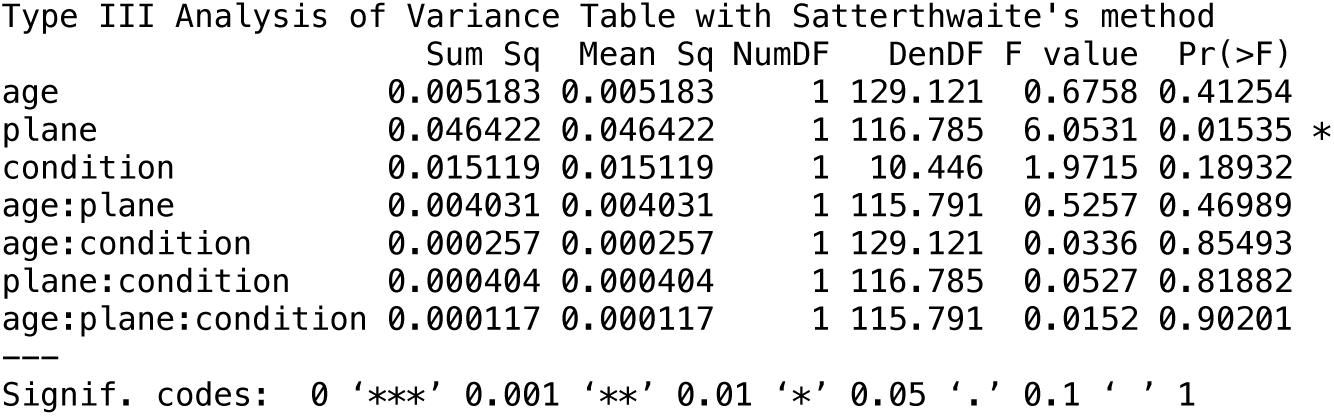

|  | Sum Sq | Mean Sq | NumDF | DenDF | F value | Pr(>F) |
| --- | --- | --- | --- | --- | --- | --- |
| age | 0.005183 | 0.005183 | 1 | 129.121 | 0.6758 | 0.41254 |
| plane | 0.046422 | 0.046422 | 1 | 116.785 | 6.0531 | 0.01535 * |
| condition | 0.015119 | 0.015119 | 1 | 10.446 | 1.9715 | 0.18932 |
| age:plane | 0.004031 | 0.004031 | 1 | 115.791 | 0.5257 | 0.46989 |
| age:condition | 0.000257 | 0.000257 | 1 | 129.121 | 0.0336 | 0.85493 |
| plane:condition | 0.000404 | 0.000404 | 1 | 116.785 | 0.0527 | 0.81882 |
| age:plane:condition | 0.000117 | 0.000117 | 1 | 115.791 | 0.0152 | 0.90201 |

Signif. codes: 0 '\*\*\*' 0.001 '\*\*' 0.01 '\*' 0.05 '.' 0.1 ' ' 1

**Table S14.**

|  | Sum Sq | Mean Sq | NumDF | DenDF | F value | Pr(>F) |  |
| --- | --- | --- | --- | --- | --- | --- | --- |
| age | 317.89 | 317.89 | 1 | 130.940 | 4.2259 | 0.0418 | * |
| plane | 53.92 | 53.92 | 1 | 121.394 | 0.7168 | 0.3989 |  |
| condition | 63.44 | 63.44 | 1 | 14.107 | 0.8433 | 0.3739 |  |
| age:plane | 19.92 | 19.92 | 1 | 120.477 | 0.2648 | 0.6078 |  |
| age:condition | 174.61 | 174.61 | 1 | 130.940 | 2.3211 | 0.1300 |  |
| plane:condition | 0.36 | 0.36 | 1 | 121.394 | 0.0048 | 0.9447 |  |
| age:plane:condition | 15.86 | 15.86 | 1 | 120.477 | 0.2109 | 0.6469 |  |

--- Signif. codes: 0 '\*\*\*' 0.001 '\*\*' 0.01 '\*' 0.05 '.' 0.1 ' ' 1

**Table S15.**
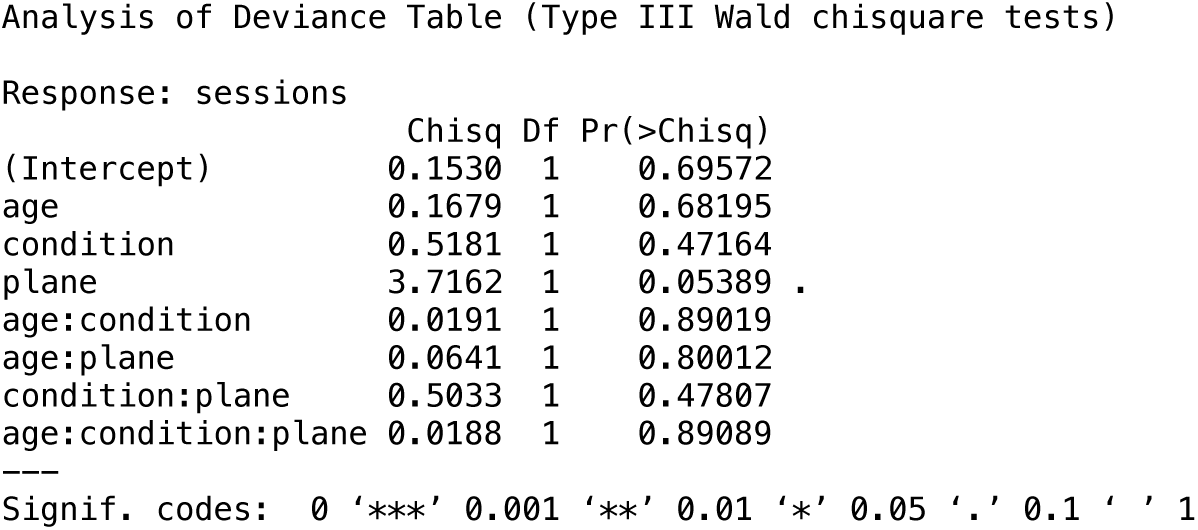

**Table S16 - related to Figure 5.**

|  | Sum Sq | Mean Sq | NumDF | DenDF | F value | Pr(>F) |
| --- | --- | --- | --- | --- | --- | --- |
| age | 2785.56 | 2785.56 | 1 | 129.719 | 16.8282 | 7.181e-05 *** |
| plane | 19.00 | 19.00 | 1 | 120.354 | 0.1148 | 0.7354 |
| condition | 154.09 | 154.09 | 1 | 13.786 | 0.9309 | 0.3513 |
| age:plane | 7.84 | 7.84 | 1 | 119.592 | 0.0474 | 0.8281 |
| age:condition | 166.36 | 166.36 | 1 | 129.719 | 1.0050 | 0.3180 |
| plane:condition | 2.22 | 2.22 | 1 | 120.354 | 0.0134 | 0.9080 |
| age:plane:condition | 349.57 | 349.57 | 1 | 119.592 | 2.1118 | 0.1488 |

| condition contrast | estimate | SE | df | t.ratio | p.value |
| --- | --- | --- | --- | --- | --- |
| c (20-23) - (24-27) | -12.72 | 3.10 | 122 | -4.100 | 0.0002 |
| uc (20-23) - (24-27) | -7.72 | 3.96 | 132 | -1.952 | 0.0796 |

Results are averaged over some or all of the levels of: plane
Degrees-of-freedom method: kenward-roger
P value adjustment: BY method for 2 tests

**Table S17 - related to1 Figure 5.**

|  | Sum Sq | Mean Sq | NumDF | DenDF | F value | Pr(>F) |
| --- | --- | --- | --- | --- | --- | --- |
| age | 11.1227 | 11.1227 | 1 | 129.302 | 12.0345 | 0.00071 *** |
| plane | 1.4302 | 1.4302 | 1 | 120.184 | 1.5474 | 0.21593 |
| condition | 0.3726 | 0.3726 | 1 | 13.856 | 0.4031 | 0.53584 |
| age:plane | 0.2790 | 0.2790 | 1 | 119.474 | 0.3018 | 0.58376 |
| age:condition | 0.1618 | 0.1618 | 1 | 129.302 | 0.1751 | 0.67635 |
| plane:condition | 0.1243 | 0.1243 | 1 | 120.184 | 0.1345 | 0.71442 |
| age:plane:condition | 3.7443 | 3.7443 | 1 | 119.474 | 4.0512 | 0.04639 * |

| condition | plane | contrast | estimate | SE | df | t.ratio | p.value |
| --- | --- | --- | --- | --- | --- | --- | --- |
| c | deep | (20-23) - (24-27) | -1.002 | 0.340 | 118 | -2.946 | 0.0323 |
| uc | deep | (20-23) - (24-27) | -0.478 | 0.411 | 129 | -1.164 | 0.6846 |
| c | sup | (20-23) - (24-27) | -0.137 | 0.305 | 123 | -0.448 | 1.0000 |
| uc | sup | (20-23) - (24-27) | -0.972 | 0.367 | 128 | -2.648 | 0.0380 |

Degrees-of-freedom method: kenward-roger P value adjustment: BY method for 4 tests

**Table S18.**

| Type III Analysis of Variance Table with Satterthwaite's method |  |  |  |  |  |  |
| --- | --- | --- | --- | --- | --- | --- |
|  | Sum Sq | Mean Sq | NumDF | DenDF | F value | Pr(>F) |
| Age | 0.193709 | 0.193709 | 1 | 119.805 | 1.6130 | 0.2065 |
| Plane | 0.176513 | 0.176513 | 1 | 112.342 | 1.4698 | 0.2279 |
| Groupe | 0.000208 | 0.000208 | 1 | 12.159 | 0.0017 | 0.9675 |
| Age:Plane | 0.260648 | 0.260648 | 1 | 110.801 | 2.1704 | 0.1435 |
| Age:Groupe | 0.090132 | 0.090132 | 1 | 119.805 | 0.7505 | 0.3880 |
| Plane:Groupe | 0.004425 | 0.004425 | 1 | 112.342 | 0.0368 | 0.8481 |
| Age:Plane:Groupe | 0.299305 | 0.299305 | 1 | 110.801 | 2.4923 | 0.1173 |

**Table S19.**
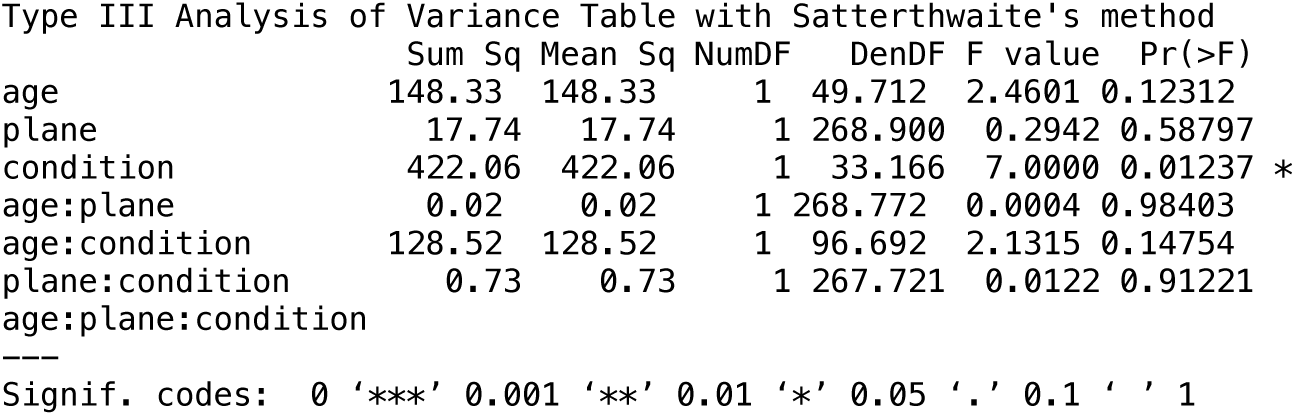

**Table S20.**
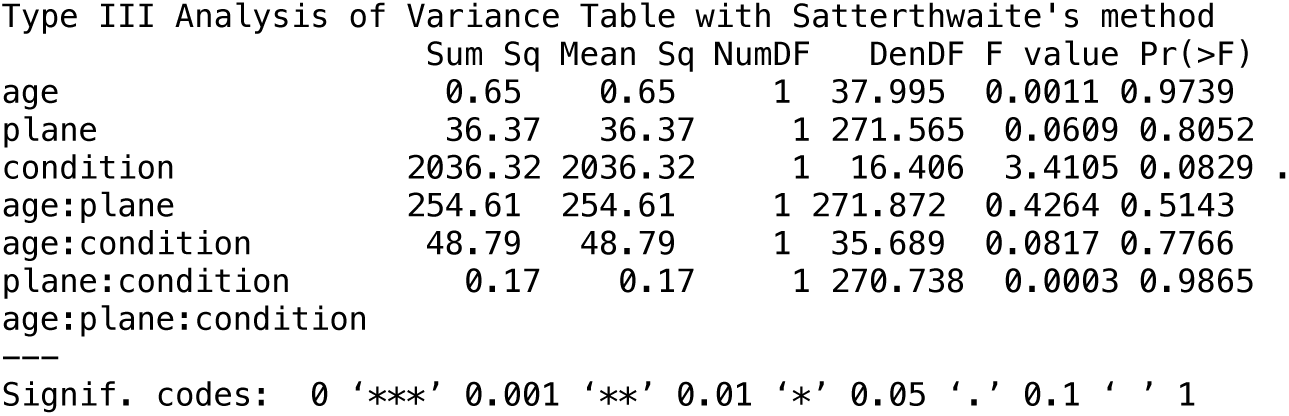

